# Single-cell RNA sequencing identifies progenitor dysfunction, inflammation and premature aging in ex vivo airway epithelium-derived from transplant recipients

**DOI:** 10.64898/2025.12.26.696372

**Authors:** Louise Bondeelle, Fedor Bezrukov, Gregory Berra, Yves Chalandon, Constant Gensous, Sheryline Loison, Federica Giannotti, Romain Messe, Jérôme Le Goff, Sophie Clément, Caroline Tapparel, Anne Bergeron

**Affiliations:** Department of Microbiology and Molecular Medicine, Faculty of Medicine, University of Geneva, Geneva, Switzerland; Pulmonology Department, Geneva University Hospitals and Faculty of Medicine, University of Geneva, Geneva, Switzerland; Department of Hematology, Geneva University Hospitals and Faculty of Medicine, University of Geneva, Geneva, Switzerland; Université Paris Cité, INSERM UMR 1342, Biology and Pathogenesis of Viral infections, Saint Louis Research Institute, F-75010 Paris, France; Virology Department, AP-HP, Hôpital Saint Louis, F-75010 Paris, France; Université Paris Cité, UMR 1153 CRESS, ECSTRRA Team, F-75010, Paris, France

**Keywords:** bronchiolitis obliterans syndrome, hematopoietic stem cell transplantation, lung transplantation, single-cell RNA sequencing, bronchial epithelium, allograft rejection

## Abstract

Pulmonary dysfunction is a common complication following hematopoietic stem cell transplantation (HSCT) or lung transplantation (LT). Bronchiolitis obliterans syndrome (BOS), an alloimmune complication of transplantation, with complex pathophysiology, contributes substantially to morbidity and mortality in these settings. We aimed to identify common epithelial features predisposing to BOS by comparing epithelia from HSCT and LT without BOS and non-transplant (NT) individuals.

We developed an ex vivo model of human airway epithelia (HAE) reconstituted from bronchial biopsies from 6 patients of each group. Using single-cell RNA sequencing, we identified two distinct epithelial profiles among transplant recipients: one resembling NT epithelium and another displaying altered cellular composition and transcriptional signatures across basal, suprabasal, club and submucosal basal duct cells, which were even more pronounced in a patient who subsequently developed BOS. This latter subgroup exhibited dysregulation of epithelial-mesenchymal transition, TNFα/NFĸB signaling and inflammatory pathways, suggesting impaired epithelial function. Moreover, these HAE demonstrated premature epithelial aging and increased expression of genes encoding damage associated molecular patterns.

Together, these findings indicate that epithelial abnormalities specific to certain transplant recipients may contribute to BOS development. Although causality cannot yet be definitively established, our data highlight the airway epithelium as a site of sustained post-transplant injury and reveal potential molecular mechanisms underlying BOS pathogenesis.

## Introduction

Pulmonary dysfunction is a common complication following hematopoietic stem cell transplantation (HSCT) or lung transplantation (LT)(1). While peri- and post-transplant events may transiently affect lung function, a subset of patients after HSCT develop a progressive, irreversible obstructive disorder referred as bronchiolitis obliterans syndrome (BOS). Similarly, after LT, some patients develop chronic allograft dysfunction (CLAD), the clinical manifestation of chronic rejection, of which BOS represents the most frequent form.

BOS is characterized histopathologically by obliterative bronchiolitis(2). In LT, BOS is defined as a persistent (>3 months) decline in forced expiratory volume in 1 second (FEV1) of ≥ 20% from the post-transplant baseline(3). In HSCT, BOS is defined by new onset of fixed obstructive ventilatory defect, (FEV1 <75% of predicted and FEV1/forced vital capacity (FVC) < 0.7), with a sustained post-transplant FEV1 decline of ≥10% proposed as an early predictor of BOS for HSCT(4). Some patients recover without progression to fixed obstruction, underscoring uncertainty in predicting BOS after FEV1 decline. BOS typically arises within three years after HSCT and five years after LT, with five-year survival rate of ≈60%(5, 6). Both chronic lung graft-versus-host disease (cGVHD) in HSCT and CLAD in LT involve alloimmune-mediated inflammation and fibrosis(1). Early diagnosis of BOS is hindered by the absence of biomarkers.

Repeated epithelial injury with impaired repair mechanisms likely triggers BOS(7, 8). Early damage may arise from HSCT conditioning regimens, surgical trauma, primary graft dysfunction (PGD) in LT, exacerbated by immunosuppressive treatment toxicity and respiratory infections. A recent American Thoracic Society (ATS) research statement underscored the need to determine the role of early epithelial changes in BOS pathophysiology(9).

The airway epithelium forms a physical and immunological barrier, with ciliated and goblet cells driving mucociliary clearance and basal, suprabasal, club, ionocytes, deuterosomal, tuft and pulmonary neuroendocrine cells contributing to immune regulation and cytokine release(10). Tissue homeostasis is maintained by airway progenitors: basal cells in the large airways and on club cells in the small airways, which also secrete Clara cell secretory protein (CCSP)(11). Suprabasal cells act as transitional progenitors derived from basal cells, differentiating into secretory or ciliated lineages. Dysfunction of these epithelial cell types likely contributes to chronic respiratory diseases(12).

Studies suggest that epithelial progenitors may also be impaired in BOS following both HSCT and LT leading to defective regeneration and airway remodeling(13). However, this remains uncertain. Basal and club cells are particularly vulnerable to immune-mediated cytotoxicity by CD8+ T lymphocytes. This epithelial damage is exacerbated by inflammatory cytokines and humoral response, resulting in impaired regeneration and progressive remodeling of the airways that is characteristic of post-transplant BOS(14).

Beyond these canonical progenitors, submucosal glands (SMGs), beneath the airway epithelial surface, may also provide an auxiliary regenerative pool(15). However, their contribution in tissue repair and their relevance to BOS pathophysiology remains incompletely understood.

Aging modulates epithelial progenitor function and influences outcomes following HSCT and LT(16, 17). In HSCT, donor-derived hematopoietic stem cells exhibit premature aging with epigenetic age acceleration, telomere attrition and chronic inflammation, increasing the risk of cGVHD(18). In LT, advanced donor age and early epithelial injury, such as PGD, accelerate airway epigenetic aging and worsen survival(17). Whether premature airway epithelial aging occurs independently after transplantation remains uncertain.

Both biological aging and transplantation-induced cellular stress promote the release of damage-associated molecular patterns (DAMPs), a hallmark of sterile inflammation and tissue aging(20) as well as in GVHD(19). These signals sustain innate immune activation, amplify alloreactivity and perpetuate a vicious cycle of defective repair and fibrosis(20, 21). Although the direct involvement of this process in BOS is not yet proven, evidence from other chronic respiratory diseases supports a pathogenic role.

In this study, we generated *ex vivo* human airway epithelium (HAE) from bronchial biopsies of HSCT and LT recipients collected within 3 years after transplantation and without BOS, alongside non-transplant patients. Using single-cell RNA-sequencing (scRNA-seq), we compared the epithelial cell proportions and cell type-specific transcriptomes. We identified two distinct transplant-derived HAE subtypes: one (T_1) resembling NT HAE and another (T_2) marked by progenitor cell composition and transcriptomic profile disruption.

Together, our findings identify the airway epithelium as a site of sustained injury after transplantation and highlight potential contributors to BO pathogenesis. Defining which transplant recipients will develop these epithelial abnormalities may enable early prediction of risk and open opportunities for targeted intervention before irreversible decline.

## Methods

### Reconstitution of human bronchial epithelia at an air-liquid interface

In our center, routine follow-up after LT includes bronchoscopy at 2 weeks, and at 1-3-6-9- and 12 months for the detection of CLAD. Both HSCT and LT have pulmonary function tests every 3 months for 1 year, then every 6 months up to 3 years. For HSCT, bronchoscopy is indicated upon ≥10% decline in FEV1 to exclude infection. NT patients underwent bronchoscopy for nodule or benign disease without prior chemotherapy or radiotherapy. All procedures were performed at Geneva University Hospitals. None of the transplant recipients met criteria for BOS. The study was approved by Swissethics N°2023-00140), all participants provided informed consent.

Three distal mucosal biopsies were collected per subject. Additional biopsies were processed as formalin-fixed paraffin-embedded (FFPE) for pathology. Respiratory samples underwent bacterial and fungal cultures, and multiplex viral PCR (Tianlong® 12-virus kit). Biopsies were excluded when any pathogen was detected.

Primary bronchial epithelial cells were expanded and differentiated at air-liquid interface (ALI) for 28 days to generate fully differentiated HAE.

### Single-cell RNA-seq sequencing

Eighteen HAE (six per group) were fixed, Hoechst stained (ThermoFisher, H1399), FACS sorted, to remove dead cells and debris. Cells were, barcoded using lipid-tagged indices, pooled and processed with the 10X Genomics Single-cell Chromium Flex Solution. Libraries were sequenced on an Illumina NovaSeq 6000 (90-cycle runs, high-output).

### Data processing and analysis

FASTQ files were processed using Cell Ranger v8.0.0 and GRCh38-2024-A human reference. Samples with ≥2,000 cells and 2,000 UMIs per cell were retained; barcode rank plots were inspected for quality. Doublets were removed using scDblFinder R package (v1.22.0). Gene expression values were normalized and log-transformed (lognorm counts).

Cell types were annotated by label transfer from the Human Lung Reference Cell Atlas v1.0(22), using scANVI(23, 24), with 1,024 variable genes for model fine-tune. Latent embeddings were visualized via UMAP.

### Differential Expression and pathway Analysis

Pseudo-bulk (total and per cell type per sample) were analyzed with DESeq2 (v1.48.2); genes with (FDR) < 0.05 were deemed significant. Gene set enrichment scores were obtained using GSVA R package (V2.0.0) with hallmark gene sets from MSigDB. To support further exploration, our data are available (https://cellxgene.cziscience.com).

## Results

### Patient characteristics

Biopsies were obtained from a total of 34 subjects, of whom 16 were excluded due to microbial contamination or absence of growth (Figure S1). Biopsies from the remaining 18 subjects (6 controls, 6 HSCT and 6 LT recipients) were used for HAE reconstitution (Figure 1A). Key demographic and clinical variables of the patients are summarized in Table 1, with additional details provided in the <u>Supplementary Material</u> *(Tables S1, S2, S3)*.

**Figure 1.**
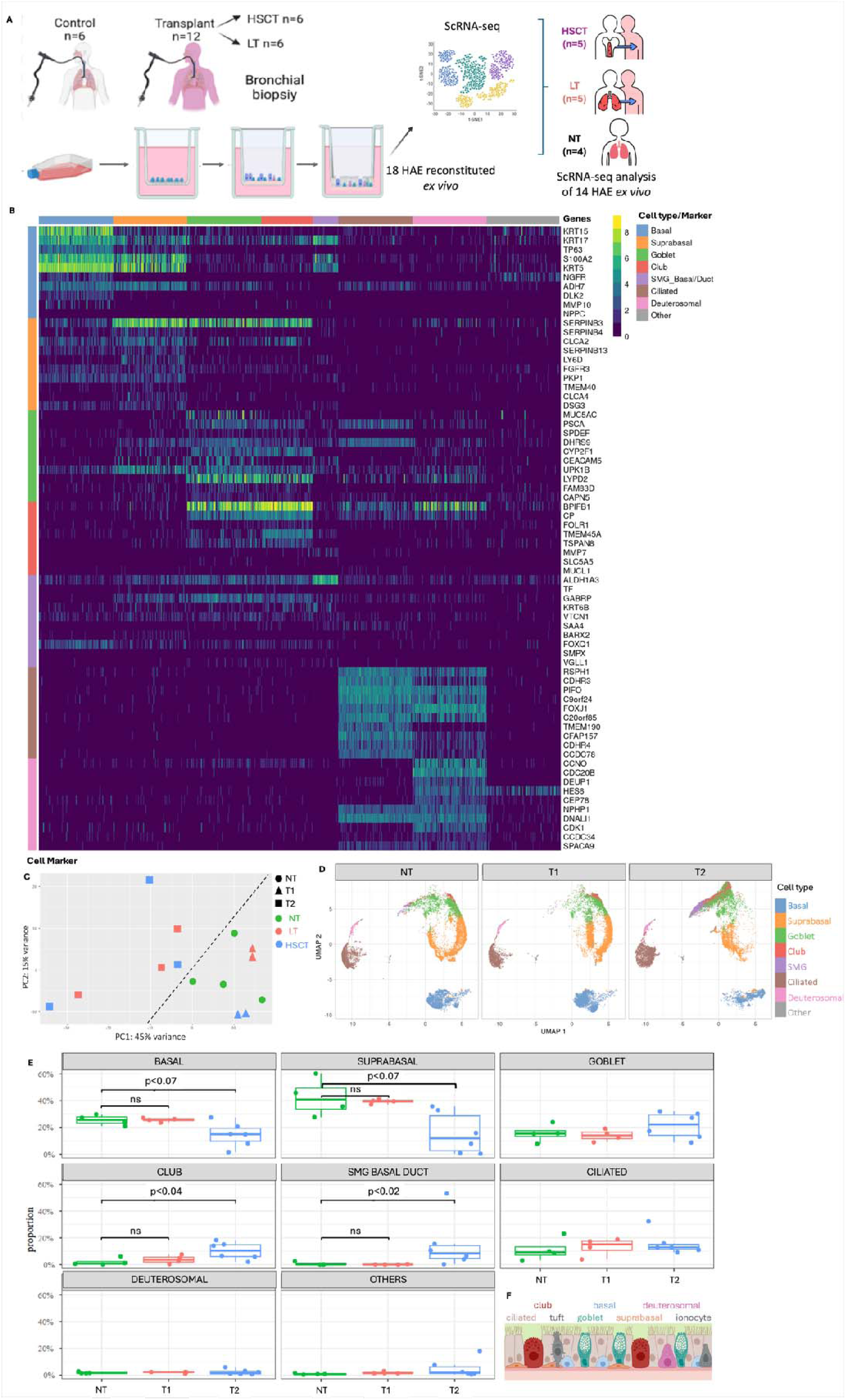
Single-cell landscape of transplant and non-transplant HAE. (A)Workflow of the study: bronchial biopsy collection, ex vivo HAE reconstitution (n = 18), followed by scRNA-seq analysis of 14 HAE samples that passed quality control. (B) Heatmap of the top ten marker genes per cell type across NT samples. (C) Principal component analysis (PCA) of total pseudo-bulk gene expression of all samples. T1 samples (triangle) and T2 samples (squares) are separated by the black dotted line, with HSCT samples shown in blue and LT samples in red. T1 samples cluster closely to NT (green circle) (D) UMAP representation of the transcriptome of all the cells in NT, T1 and T2 groups. Cells are colored according to their type (E) Cell type proportions per individual across groups. The horizontal bar indicates the median. Each dot represents an individual: NT (green) and T1 (25), T2 (pale blue). (F) Representative airway epithelial cell populations.

**Table 1.**
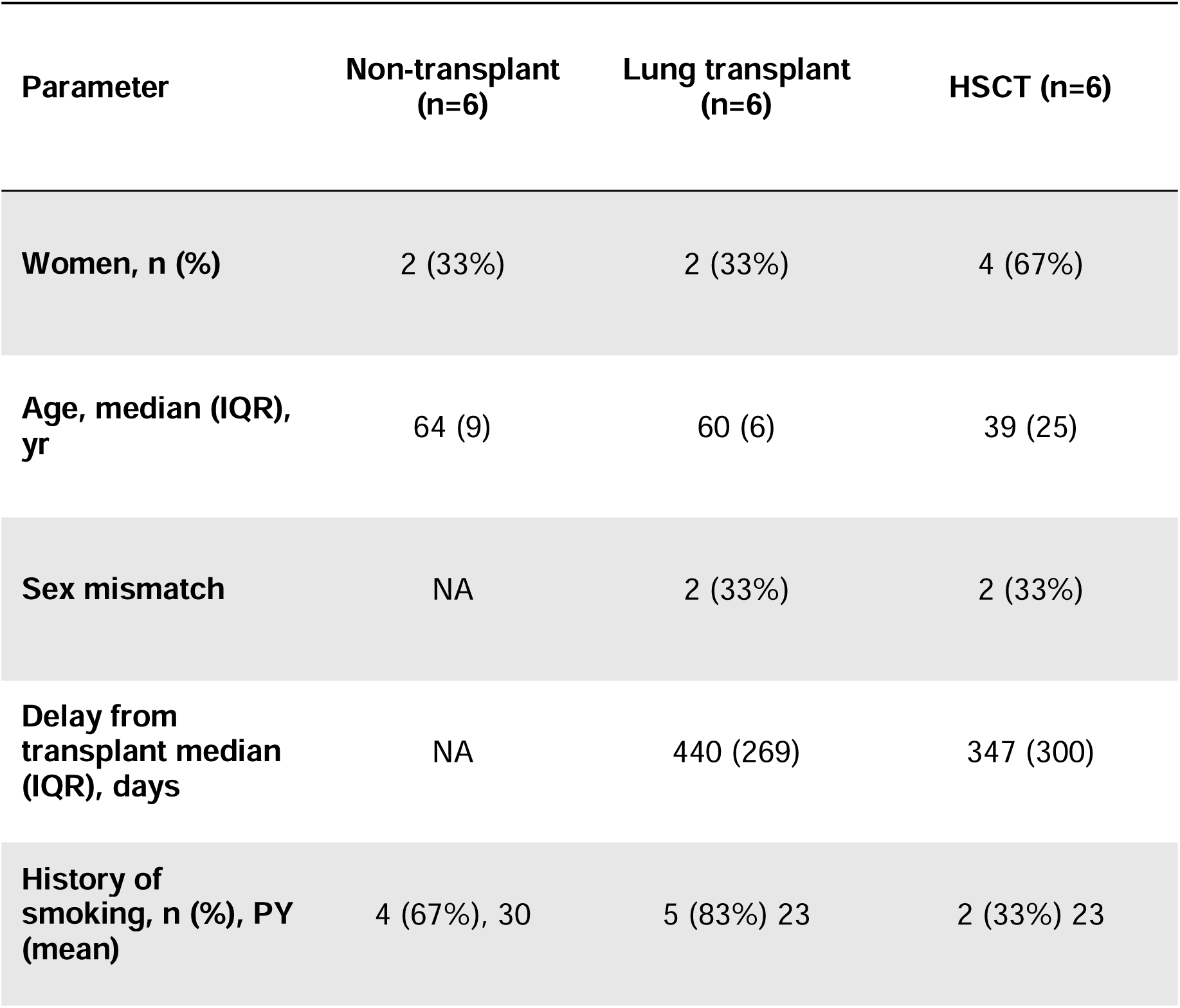
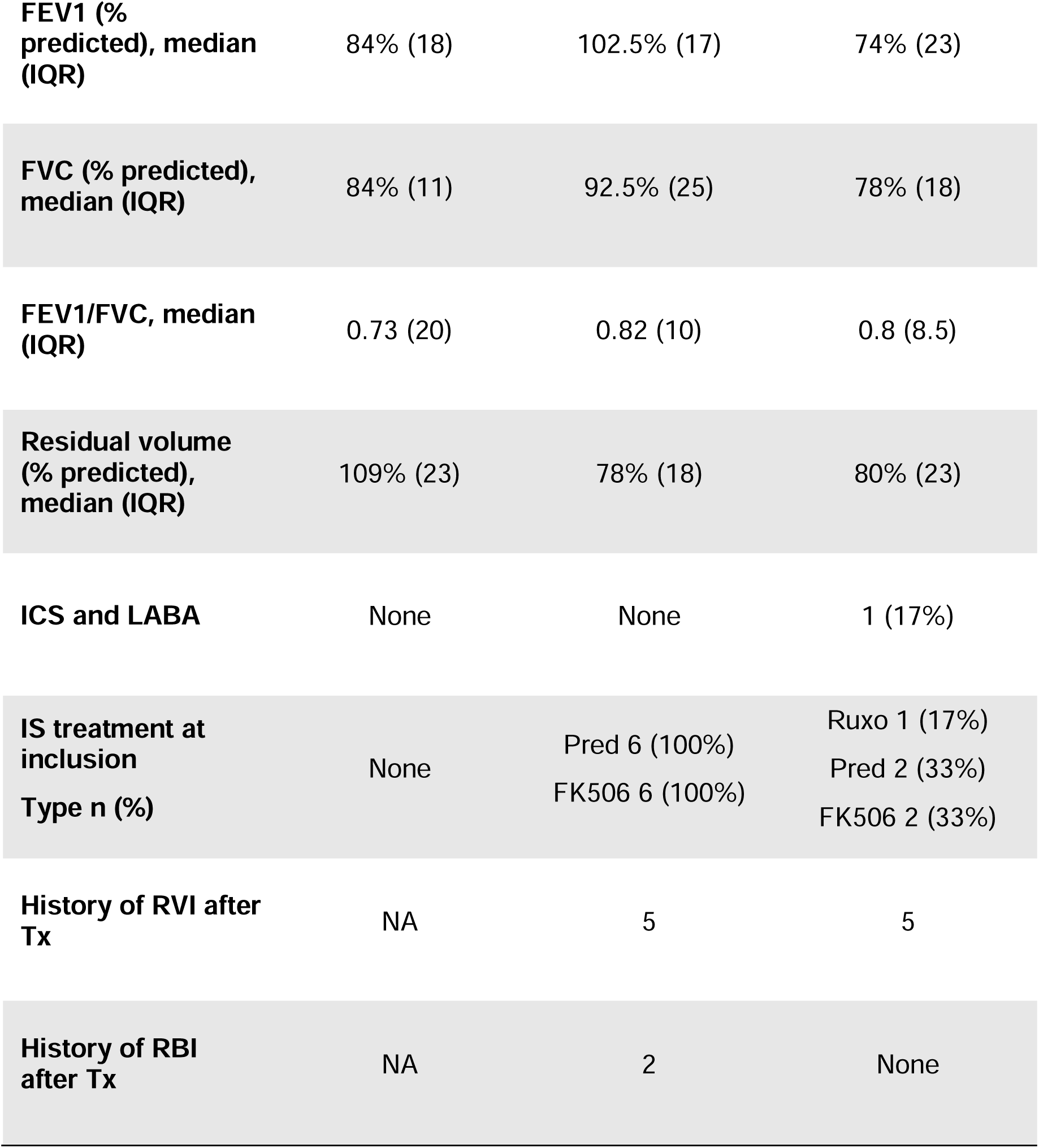
Patient characteristics IQR: interquartile (Q1-Q3), yr: year. PY: pack-year (number of cigarette packs per day x the number of years of smoking) FEV1: forced expiratory volume in 1 second, FVC: forced vital capacity, RV: residual volume, ICS: inhaled corticosteroid, LABA: long-acting beta2 agonists, IS: immunosuppressive, Pred: prednisone; Ruxo: Ruxolitinib; FK: tacrolimus, RBI: respiratory viral infection, RVI; respiratory viral infection, NA: not available.

### Single-cell RNA sequencing reveals different transcriptomic profiles between transplant and non-transplant HAE and highlights the presence of two transplant subclusters

To delineate the cellular landscape of reconstituted HAE, we performed scRNA-seq across all samples. Four HAE were excluded due to insufficient sequencing quality (Figure S1), leaving 14 HAE (4 NT, 5 LT and 5 HSCT) for downstream analysis (Figure 1A). Following standard quality control and processing, a total of 76,109 cells (NT =□17,742, LT = 32,367, HSCT = 26,000) were retained for analysis. Cell types were inferred from the mRNA expression profiles by comparison with annotated LungMAP atlas (label transfer), identifying eight major distinct cell types corresponding to canonical bronchial epithelial lineages, confirmed by established marker genes (Figure 1B). The Principal Component Analysis (PCA) of the average all cell combined transcriptome (bulk) identifies three major groups (Figure 1C): the NT group and two transplant-associated clusters (T1 and T2). The T1 cluster (LT4, LT5, HSCT4 and HSCT6) exhibited proximity to the NT samples, whereas the T2 cluster (LT1, LT3, LT6, HSCT2, HSCT3 and HSCT5), clearly segregated from NT. Subsequent analyses focused on T1, T2 and NT comparisons. UMAP projections confirmed that T1 shared more similarities in transcriptional behavior to NT than to T2 (Figure 1D). Analysis of cell-type proportions revealed moderate intergroup and interindividual variability (Figure 1E). Compared to T2, NT HAE tended to be enriched for basal (26.2%) and suprabasal cells (38.5%) and contained significantly low fractions of club (2.3%) and SMG cells (0.8%). T1 mirrored NT in basal (25.7%), suprabasal (39.6%) and SMG cells (0.2%) with a slight increase in club (4.3%). In contrast, T2 tended to display a depletion of basal (16.1%) and suprabasal cells (15.6%) accompanied by a substantial and expansion of club (11.2%) and SMG cells (9.7%) (Figure 1E). A schematic summary of airway epithelial organization and cell subsets is provided in Figure 1F. We next compared the transcriptomic profiles of the different cell populations.

### ScRNA-seq analysis identified a distinct transcriptomic signature in basal cells from transplant recipients of the T2 cluster

Basal cell, critical lung stem cells implicated in BO(26), were analyzed in detail due to their altered proportion in T2 *versus* NT and T1. A total of 16,733 basal cells were analyzed: 4,644 NT cells (26.2%), 7,191 T1□cells (25.7%) and 4,898 T2 cells (16.1%), with individual variability illustrated in Figure 2A.

**Figure 2.**
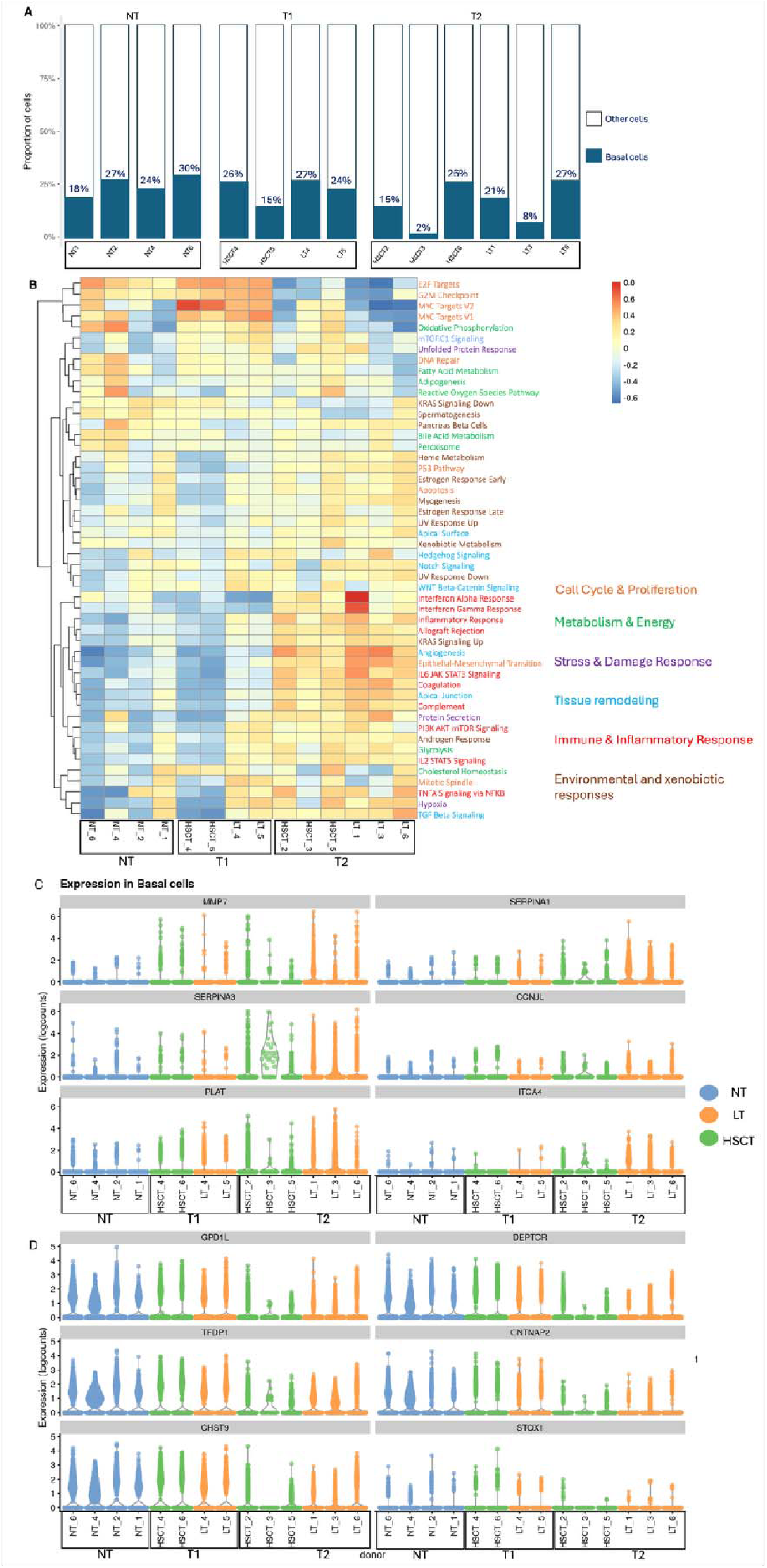
Transcriptomic profile of basal cells across all groups (A) basal cells proportion across all samples **(B)** Heatmap of gene set enrichment scores of 50 signaling pathway gene expression across samples, with subgroups indicated (NT: non-transplant; T1: transplant group 1; T2: transplant group 2). These pathways correspond to six biological processes: cell cycle and proliferation, metabolism and energy (green), stress & damage response (purple), tissue remodeling (blue), immune and inflammatory responses (red) and environmental & xenobiotic responses (brown). (C) Violin plots of top upregulated genes in basal cells of T2 and T1 compared to NT (D) Violin plots of top downregulated genes in basal cells of T2 and T1 compared to NT. In C and D, adjusted p value (padj) for the comparison between NT and T2 ranged from 1.2 10e-23 to 2.1 10e-7. For the comparison of T1 and T2, padj value raged from 3.1 1oe-13 to 1.1 10e-2. In contrast, comparison between NT and T1 showed no significant differences, padj >0.05.

Basal cells from T2, compared to NT and T1, showed upregulation of pathways essential for airway epithelial function (Figure 2B). The six top upregulated genes were MMP7 (extracellular matrix remodeling), SERPINA1 and SERPINA3 (viral maturation and proteinase inhibitor), CCNJL (cell adhesion regulation), ITGA4 (cell adhesion receptor mediating immune cell recruitment) and PLAT (fibrin degradation and tissue remodeling) (Figure 2C). Additional upregulated genes (Table S4) were linked to hypoxia, KRAS signaling, inflammation, glycolysis, coagulation, estrogen response and allograft rejection.

Conversely, the six most downregulated genes in T2 included GPD1L (hypoxia and oxidative stress responses), DEPTOR (mTOR pathway inhibitor controlling cell growth and autophagy), TFDP1 (cell cycle and DNA replication regulation), CNTNAP2 (cell adhesion regulation), CHST9 (extracellular matrix regulation) and STOX1 (oxidative stress and differentiation programs regulation) (Figure 2D), alongside reduced expression of SOX2 and SOX21 (progenitor regulation); FOXN1 (immune development) and CLDN8 (tight junction) (table S4).

These findings suggest that basal cells in T2 adopt a stressed, pro-fibrotic phenotype with epithelial-mesenchymal transition (EMT), inflammatory and metabolic stress pathways, alongside downregulation of epithelial integrity and immune genes, reflecting impaired repair and a shift towards a stress- and inflammation-prone state.

In T2 samples, suprabasal cells, normally intermediate progenitors between basal and differentiated lineages, exhibited a transcriptomic profile closely resembling basal cells including a pro-fibrotic program with increased EMT, inflammatory and metabolic stress, alongside reduced expression of genes involved in epithelial integrity. Detailed results for other cell types are provided in the Supplementary data (Fig S2-S7).

### Single-cell transcriptomic analysis revealed two distinct club cell subpopulations in HAE with distinct proportions across conditions

Club cells, another key progenitor population implicated in BO, were significantly more abundant in T2 (Figure 1E). A total of 5,020 club cells were analyzed, with 400 (2.3%), 1,197 (4.3%) and 3,423 (11.2%) cells in NT, T1 and T2 groups respectively (Figure 1E).

Club cells were identified by SCGB1A1 (CCSP) expression and segregated into two transcriptionally distinct subtypes based on 20 markers genes (figure 3A).

**Figure 3.**
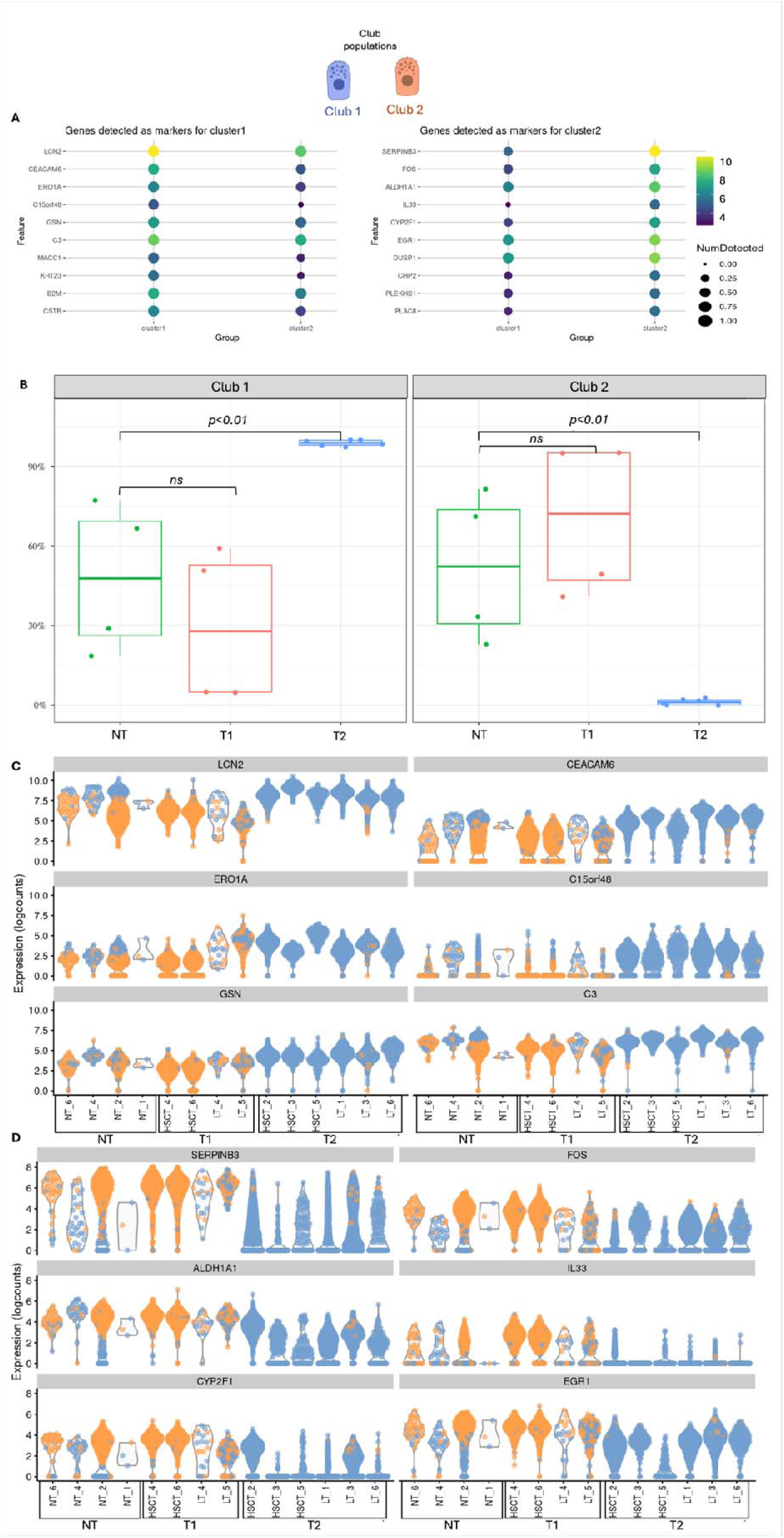
Subpopulations of Club cells (A) Expression patterns club cell markers across club1 and club2 subgroups. Throughout the Figure, club 1 is shown in blue and club 2 in orange. (B) Proportion of club 1 and club 2 subclusters across groups, each dot is a sample. The NT and T1 groups display an even distribution, whereas the T2 group is enriched for club 1 (99%) (C-D) Violin plots showing the top 6 upregulated genes in club 1 across all groups (D) and top 6 upregulated genes in club 2(E). In C and D, adjusted p value (padj) for the comparison between NT and T2 ranged from 1.2 10e-7 to 2.2 10e-3. For the comparison of T1 and T2, padj value raged from 2.1 1oe-6 to 1.1 10e-5. In contrast, comparison between NT and T1 showed no significant differences, padj >0.05.

Club 1 was defined by expression of LCN2 (epithelial stress), CEACAM6 (inflammatory adhesion), ERO1A and C15orf48 (oxidative stress) GSN (cytoskeletal organization) and C3(complement activation), MACC1 (remodeling), KRT23 and CSTB (stress response) and B2M (immune activation). These markers indicate a pro-inflammatory, stress-associated club cell state with limited regenerative capacity. In contrast, Club 2 was characterized by expression of SERPINB3 (epithelial protection), FOS; PLEKHS1, PLAC8 and EGR1 (proliferation/repair), ALDH1A1, EGR1 (epithelial regeneration) and IL33 (immune signaling), CYP2F1 and CHP2 (metabolism), DUSP1 (anti-inflammatory) (figure 3A). These markers indicate a regenerative club cell state.

Club1 and Club2 were evenly distributed in T1 and NT, whereas T2 was dominated by Club1, with a near-complete loss of Club 2, indicating a shift toward a pro-inflammatory and poorly regenerative state (figure 3B). The individual expression levels of the top six markers defining each club cell subpopulation across individual samples are shown in Figures 3C and D.

### Selective presence and distinct pathways signature of SMG basal duct cells in T2

SMG basal duct cells, which line the duct of submucosal gland (Figure 4A) were detected predominantly in T2, and at very low levels in NT and T1. A total of 5426 cells were identified: 473 (0.8%) in NT, 140 (0.5%) in T1 and 4813 (15.8%) in T2, *p=0.02* (Figure 1D). SMG accounted for 53.2% of all analyzed cells in HSCT3 and were absent in LT5 (figure 4B). Given the limited number of SMG cells, pathway analysis only indicates general trends (Fig S7). In T2, SMG cells exhibited modest increases in pathways related to inflammation, tissue remodeling and epithelial stress compared to NT and T1. The selective presence of SMG basal duct cells in T2 together with their stress-associated pathway patterns, suggest that these cells in some transplant airways may adopt a more basal-like, repair-oriented state.

**Figure 4.**
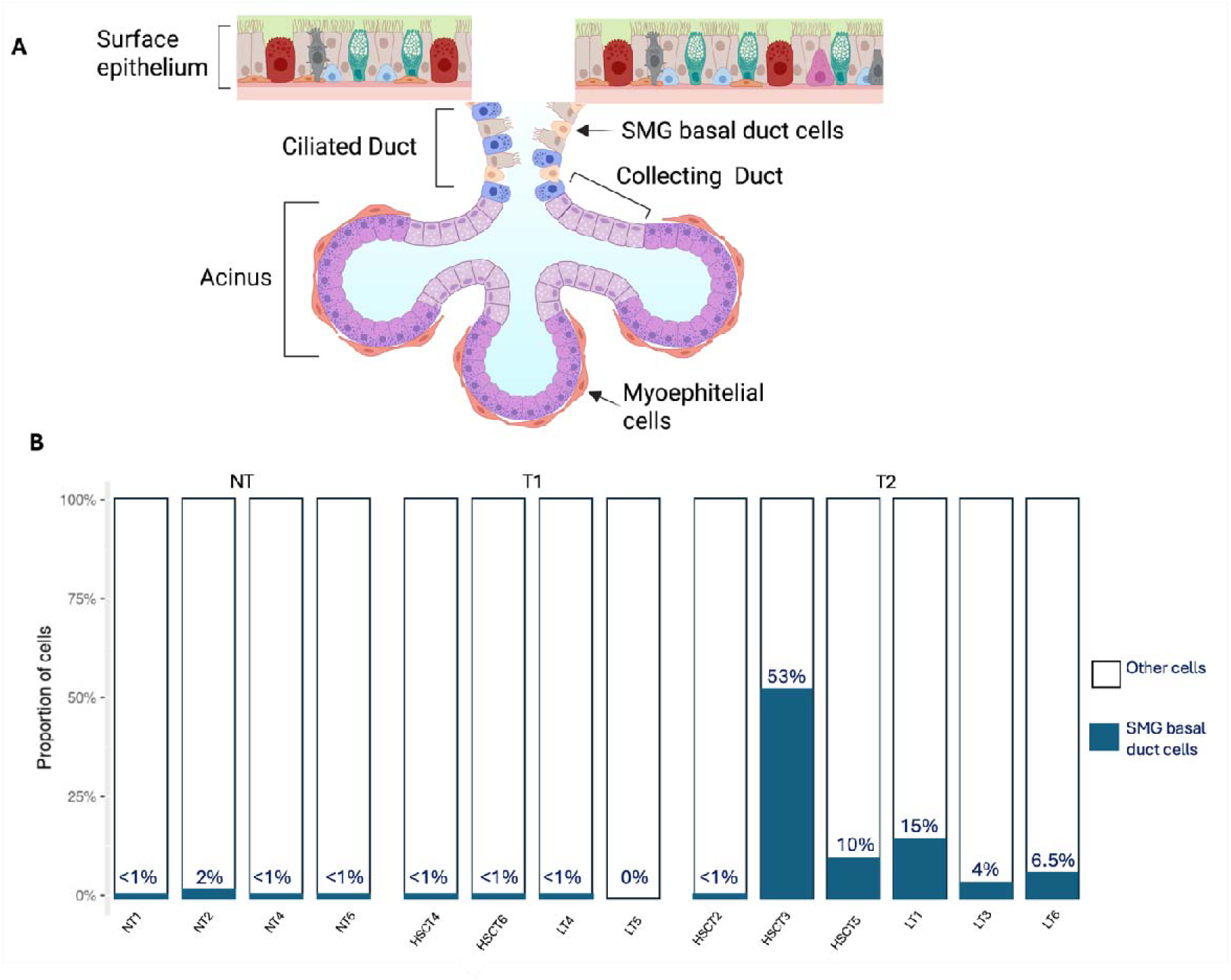
SMG basal duct cells in T1, T2 and non-transplant HAE. (A) Structure of submucosal gland. (B) SMG basal cells proportion across all samples

### ScRNA-seq revealed premature aging signatures in HAE from T2 recipients

Because accelerated aging has been proposed as a feature of bronchial epithelial cells following transplantation, potentially contributing to impaired regeneration, stem cell exhaustion, immune dysregulation and ultimately BOS, we investigated aging-related genes expression in HAE(18, 27). This framework encompasses canonical hallmarks of aging, including genomic instability, telomere attrition, epigenetic alterations, loss of proteostasis, deregulated nutrient sensing, mitochondrial dysfunction, cellular senescence and impaired intercellular communication(28).

Transcriptomic analysis identified 35 aging-related genes differentially expressed in T2 HAE compared to NT, predominantly in basal and supra-basal cells (Figure 5A). Well-established aging markers exhibited consistent changes across these cells, including upregulation of CDKN2A, CDKN2B, SERPINE1, CDKNB2, CSNK1E, IRS2, SQSTM1, IGF1R, BAK1, NOG, PDGFB, APOE and PLAU, and downregulation of PIK3R1, TP73, FOS, XPA, EPS8, HSPA1A and LMNB1. Some changes were cell type-specific, including reduced PTEN in deuterosomal cells, upregulation of FOXO3 and IRS2 in ciliated cells, and downregulation of LMNB1 and BDNF in goblet cells (Figure 5A).

**Figure 5.**
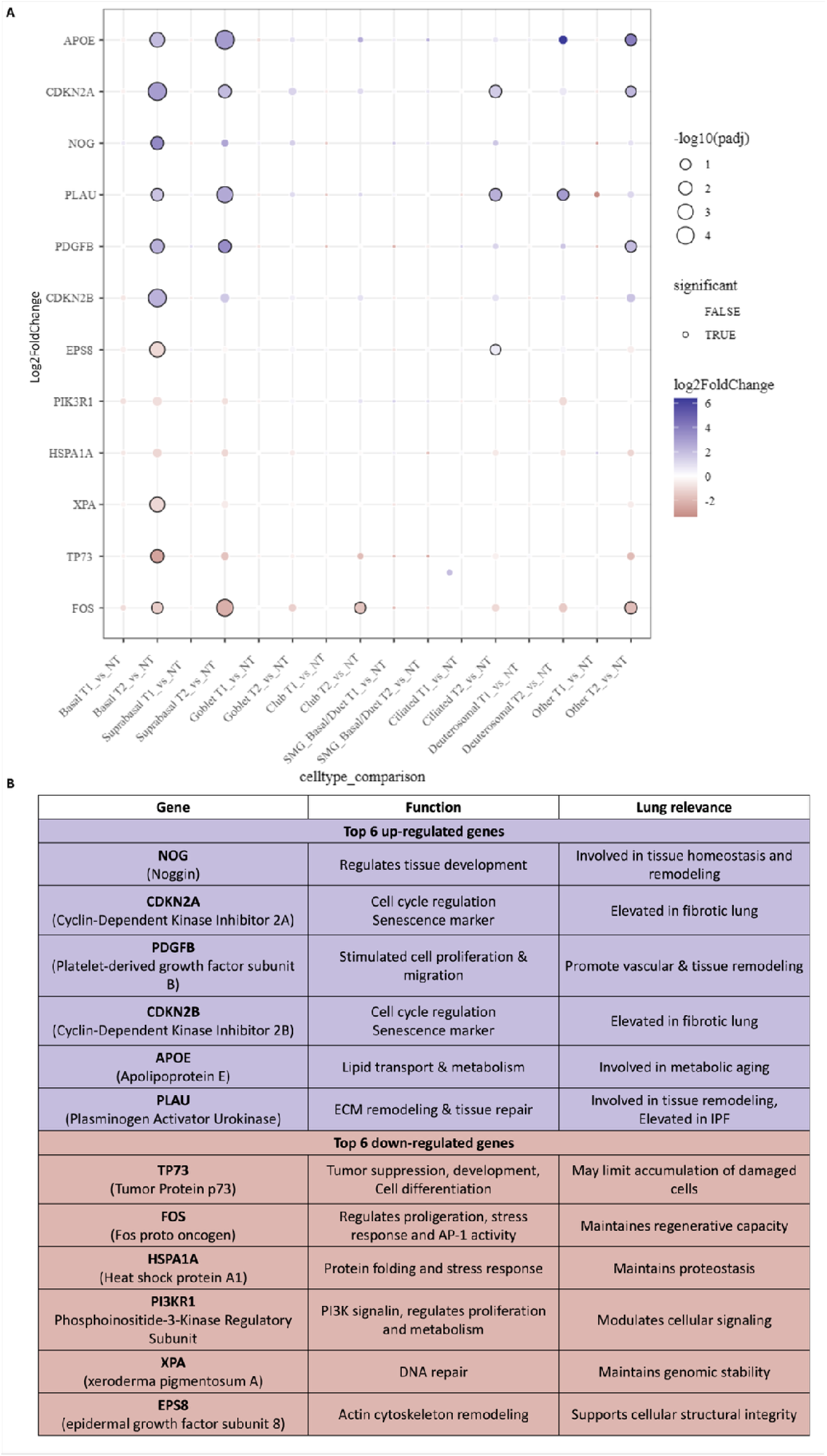
Aging-related genes differential expressed between T1, T2 and non-transplant HAE. (A) Dot plots of the top 12 genes out of 35 genes, 6 upregulated and 6 downregulated (from 307 genes in GenAge database) differentially expressed in at least one cell type between transplant versus non-transplant HAE. Gene expression was compared between T1 and NT or between T2 and NT for each cell type. Color scale represents log_2_FC (blue-violet, up to +4; orange down to -2); dot size reflects significance (larger = lower adjusted p values). (B) Functions and lung relevance of the top six up- and downregulated genes known as implicated in aging pathways.

The six most up. and down-regulated genes (Figure 5B) were enriched in HSCT3, HSCT5 (30-year-old), and LT1, LT3 and LT6 (from 55 to 70-year-old) (fig. S7), whereas the oldest NT individual, NT6 (76 years) showed lower expression. These results indicate widespread and cell type-restricted transcriptional alterations consistent with accelerated bronchial epithelial aging following transplantation. Complete gene lists are in Table S5.

### Dysregulation of DAMPs-linked genes in T2 HAE

Given the role of DAMPs in post-transplant tissue injury, we investigated their expression in HAE. Thirteen DAMPs-associated genes were significantly upregulated and three downregulated in T2, predominantly in basal and suprabasal cells (Figure 6A). The seven most strongly upregulated genes were **IL1A** (alarmin, pro-inflammatory cytokine), **FN1** and **LAMA1** (extracellular matrix glycoproteins), **VCAN** (matrix proteoglycan), **CYP26B1** (enzyme of retinoic acid degradation), **LGALS1** (extracellular matrix glycoproteins) and **HAS3** (matrix glycosaminoglycan). Among downregulated genes, **IL33** (Alarmin), was decreased in ciliated cells of T2 *vs* NT. In contrast, **CIRBP** (stress-induced RNA-binding protein) and **S100A8** (inflammatory protein) were reduced in rare cell types and ciliated cells of T1 *vs* NT, respectively. No significant changes were observed in club, goblet or SMG cells. Figure 6B summarizes their functional roles in epithelial injury and repair. Complete gene lists are provided in Table S5.

**Figure 6.**
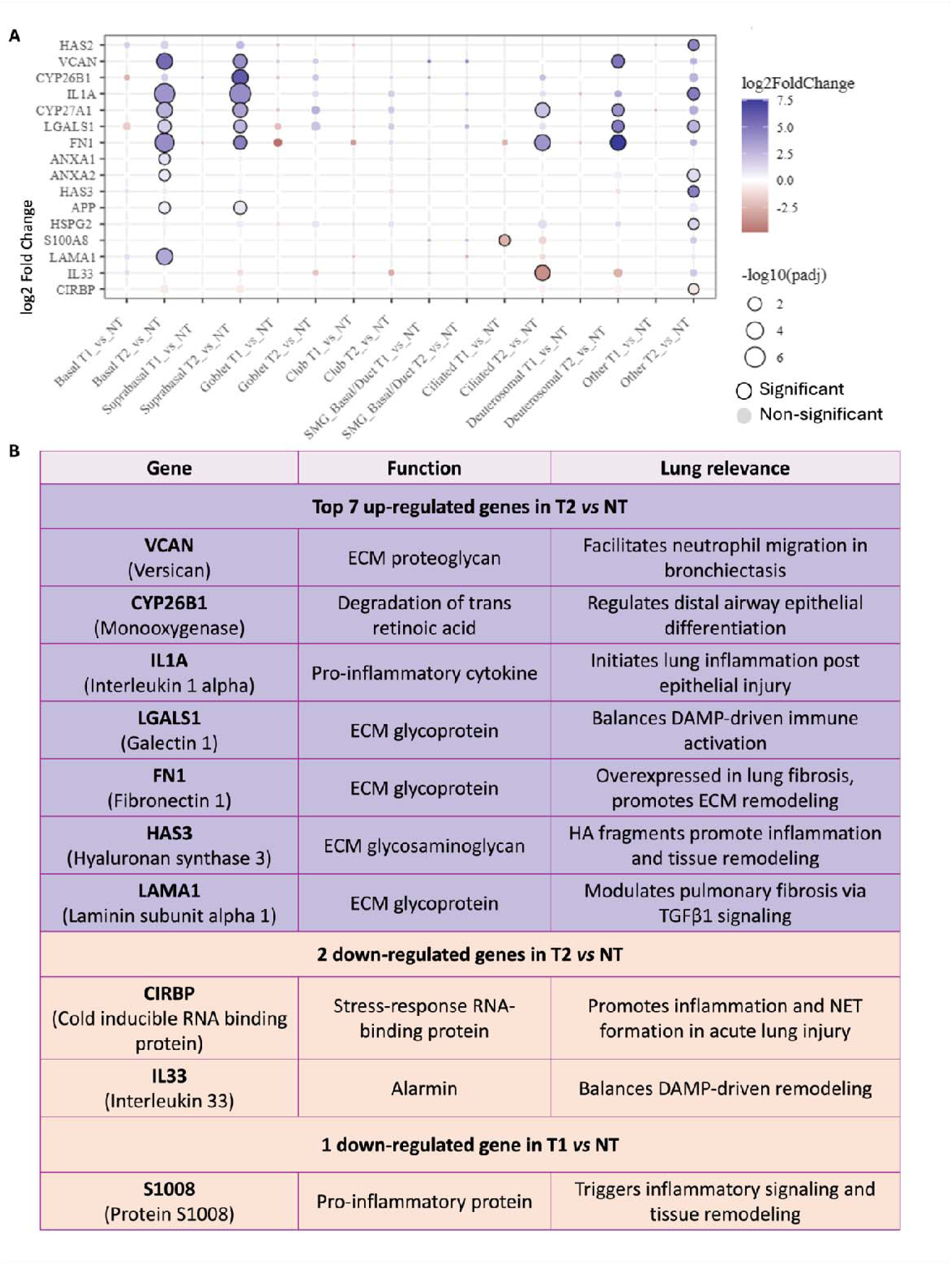
DAMPs-related genes differentially expressed between T1, T2 and non-transplant HAE. (A) Dot plots showing 16 DAMP-associated genes differentially expressed in at least one cell type between transplant and non-transplant HAE. For each cell type, gene expression was compared between T1 vs NT and between T2 vs NT. Color scale represents log_2_FC (blue violet, up to +4; orange down to -2); Dot size reflects significance (larger = lower adjusted p values). (B) Summary of functional and lung-specific relevance of the top seven upregulated genes and three down-regulated genes.

### Distinct epithelial signature in an outlier T2 sample preceding BOS onset

One T2 HAE sample, HSCT3, was identified as an outlier and analyzed separately in relation to clinical follow-up. HSCT3 displayed a distinct transcriptional and cellular profile, including increased abundance of SMG basal duct cells (Figures 7A-B). In basal cells, 10 genes were most strongly dysregulated compared with NT and the rest of T2, including upregulation of DAMPs- and aging-related genes (Figure 7C-D). Longitudinal PFT follow-up revealed that HSCT3 was the only case progressing to BOS, occurring 3 months after inclusion (Figure 7E).

**Figure 7.**
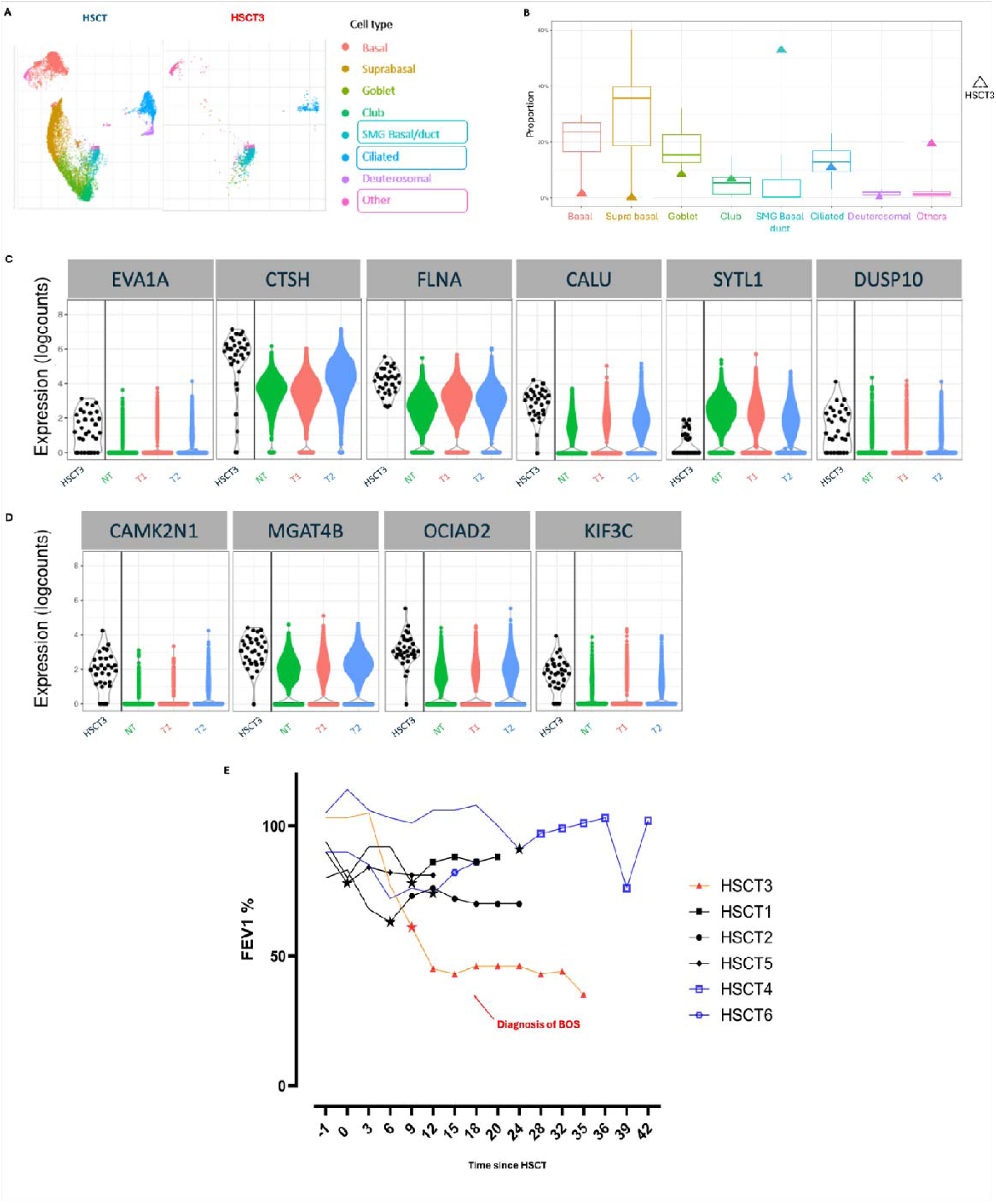
Functional and transcriptomic characteristics of the HSCT outlier (A) UMAP of scRNA-seq showing poorly differentiated epithelium in HSCT3 versus other HSCT samples; (B) Cell type proportion in HSCT3 (triangle) compared with all HAE (box plot) with the mean represented by the line; (C) Upregulation of DAMPs-related genes and (D) Aging-related genes; padj values ranged from 6.2 10e-5 to 2.9 10e-2 (E) FEV1 trajectories of all HSCT before (continuous line), at the time of biopsy (stars) and after biopsy (points), showing persistent decline and BOS development in HSCT3 (red); HSCT from T1 subgroup are in blue and HSCT from T2 subgroup are in black.

### Clinical post-transplant injury burden parallels ex vivo epithelial phenotypes

Clinical data (Tables S1-3) broadly paralleled *ex vivo* observations. T1 recipients showed a low post-transplant injury burden with preserved lung function and few respiratory infections; both LT had non-smokers donors. Among HSCT, one experienced limited mild cGVHD. In contrast, T2 recipients experienced a higher injury burden. All HSCT recipients had recurrent RVI and had a history of mild to severe extra-thoracic cGVHD. Two LT donors were smokers, three LT had severe gastroesophageal reflux, and one LT had prolonged ischemic time. None of the LT had acute allograft dysfunction. No acute allograft dysfunction occurred in either group of LT.

## Discussion

Our findings confirm that *ex vivo* HAE from bronchial biopsies of transplant recipients without BOS (HSCT and LT) differ from NT controls, and segregate into two distinct transplant phenotypes: T1 and T2. Single-cell profiling revealed that T1 closely mirrored NT epithelium, whereas T2 exhibited four distinctive features: (i) altered progenitor cell biology, with dysfunction of basal and club cells and expansion of SMG progenitors; (ii) a shared pro-inflammatory transcriptional reprogramming marked by EMT, IL2/STAT5 and TNFα/NFĸB activation; (iii) premature epithelial aging independent of chronological age; (iv) sustained DAMP signaling.

Altered progenitor biology emerged as a central feature of T2. Basal cells, essential for epithelial maintenance and repair(29), exhibited transcriptional exhaustion with impaired renewal capacity within a pro-inflammatory environment. Club cells, which combine secretory and progenitor functions, were profoundly remodeled, with depletion of a canonical reparative subset and enrichment of a pro-inflammatory subset in T2(30). This mirrors experimental models where club cell depletion promotes BO in both LT(31) and HSCT(32), and clinical observations linking reduced CCSP levels and BO risk(33). Even in clinically stable T2 recipients, this shift likely compromises epithelial repair, mucosal defense and immunomodulation.

In parallel, SMG-derived progenitors, a niche involved in mucosal defense(34), were expanded in T2 HAE but nearly absent in T1, suggesting recruitment of reserve progenitors to compensate exhaustion of canonical basal and club lineages. Although SMG hypertrophy characterizes COPD but not idiopathic pulmonary fibrosis, the underlying disease in several LT recipients(35), its presence in both HSCT and LT HAE indicates that transplantation itself reshapes progenitor hierarchies. The predominance of SMG basal duct cells in the HSCT who later developed BOS suggests that abnormal reliance on reserve progenitors may increase disease susceptibility.

A shared pan-epithelial transcriptional program distinguished T2 from T1 and NT. Across basal, suprabasal, ciliated, goblet and deuterosomal, T2 exhibited concordant activation of EMT, IL6/JAK/STAT3, TGFβ, hypoxia, TNFα/ NFĸB and allograft rejection. These global reprogramming rather than isolated cell-type-specific changes, was consistent across all LT and HSCT. Enhanced of JAK-STAT signaling in basal cells aligns with reports linking enhanced MHC-I expression and cytotoxic T-cell-mediated injury in CLAD(36), while TGFβ activation reinforces role in fibrotic airway remodeling, compounded by loss of reparative club cells(37).

Accelerated epithelial aging was another striking feature. Signatures of telomere attrition, mitochondrial dysfunction, senescence and genomic instability were enriched in T2 but not T1, with the strongest signals observed in young HSCT recipients and in the individual who later developed BOS. While premature aging has been reported in the hematopoietic and immune compartment following HSCT and in lung parenchyma following LT and also in pulmonar fibrosis, this provides to our knowledge, the first evidence of airway epithelial premature aging in HSCT(16, 38, 39). Our data suggest transplantation accelerates epithelial aging independently of chronological age, impairing regenerative fitness and amplifying chronic inflammation, thereby predisposing to BOS.

Consistent with this concept, DAMPs-related signaling, a hallmark of sterile injury and aging-associated inflammation, was upregulated in T2. DAMPs activate innate immune receptors(40), sustain inflammation, potentiate alloimmune responses in LT and amplify cGVHD in HSCT(21). DAMPs-associated transcripts (FN1, IL6, STAT3 and TGFβ1), were enriched particularly in basal and suprabasal cells in T2, whereas T1 exhibited reduced expression of the pro-inflammatory DAMP S1008. Clinical outcomes mirrored the *ex vivo* data. T1 recipients had largely uncomplicated post-transplant courses with few infections, supporting epithelial recovery and preserved lung function. In contrast, T2 recipients experienced recurrent viral infections and additional injuries (reflux, prolonged ischemic time), consistent with greater epithelial damage and altered transcriptional profiles. These findings suggest that cumulative epithelial injury, rather than a single insult, drives divergent outcomes, with even clinically stable T2 recipients showing epithelial priming toward immune activation and maladaptive remodeling, potentially predisposing to BOS.

**Study limitations** include modest sample size and differences in clinical trajectories between LT and HSCT. Although T2 LT recipients had not yet developed functional decline, T2 HSCT recipients already showed FEV1 reduction predicative of BOS. The “steady state” designation in HSCT is therefore imperfect, though most recipients recovered lung function. Despite limited immune profiling, scRNA-seq revealed reproducible epithelial altercations that nay precede overt disease.

**In conclusion**, this single-cell atlas of transplant-derived airway epithelium, identifies convergent epithelial maladaptation characterized by reduced regenerative capacity, inflammatory club and basal cells, SMG progenitor expansion, premature aging and DAMP signaling. These features represent candidate biomarkers and therapeutic targets and underscore the need to identify predictive factors of epithelial dysregulation, including epigenetic alterations.

## Ethics approval

The project received the approval of the ets Ethics Committee. Project number 2023-00140 approved on March 7^th^ 2023.

## Authorship Contributions

LB, AB and CT wrote the manuscript. FB made the bioinformatic analysis. LB, FB, GB, YC, CG, SL, FG, RM, JLG, SC, CT and AB revised it critically for important intellectual content. All authors approved the final version of the manuscript; moreover, all authors agree to be accountable for all aspects of the work in ensuring that questions related to the accuracy or integrity of any part of the work are appropriately investigated and resolved.

## Conflict of Interest Disclosures

The authors declare that they have no relevant conflicts of interest.

Y.C. has received consulting fees for advisory board from MSD, Novartis, Incyte, BMS, Pfizer, Abbvie, Roche, Jazz, Gilead, Amgen, Astra-Zeneca, Servier, Takeda, Pierre Fabre, Medac; Travel support from MSD, Roche, Novartis, Pfizer, BMS, Gilead, Amgen, Incyte, Abbvie, Janssen, Astra-Zeneca, Jazz, Pierre Fabre, Sanofi all via the institution.

## Acknowledgments

This article is based on a study first reported in a previous manuscript.

The authors thank the FACS platform of the University of Geneva for technical assistance with flow cytometry and the iGE3 Genomics Platform (UNIGE) for technical help.

This work was supported by the Fondation Privée des Hôpitaux Universitaires de Genève (HUG), the Fondation La Laurène, Ben ficheur and ORGANOVIR.

## Notes

### Competing Interest Statement

The authors have declared no competing interest.

