## supplementary materials for "Single-cell RNA sequencing identifies progenitor dysfunction, inflammation and premature aging in ex vivo airway epithelium-derived from transplant recipients"

** co last authors*

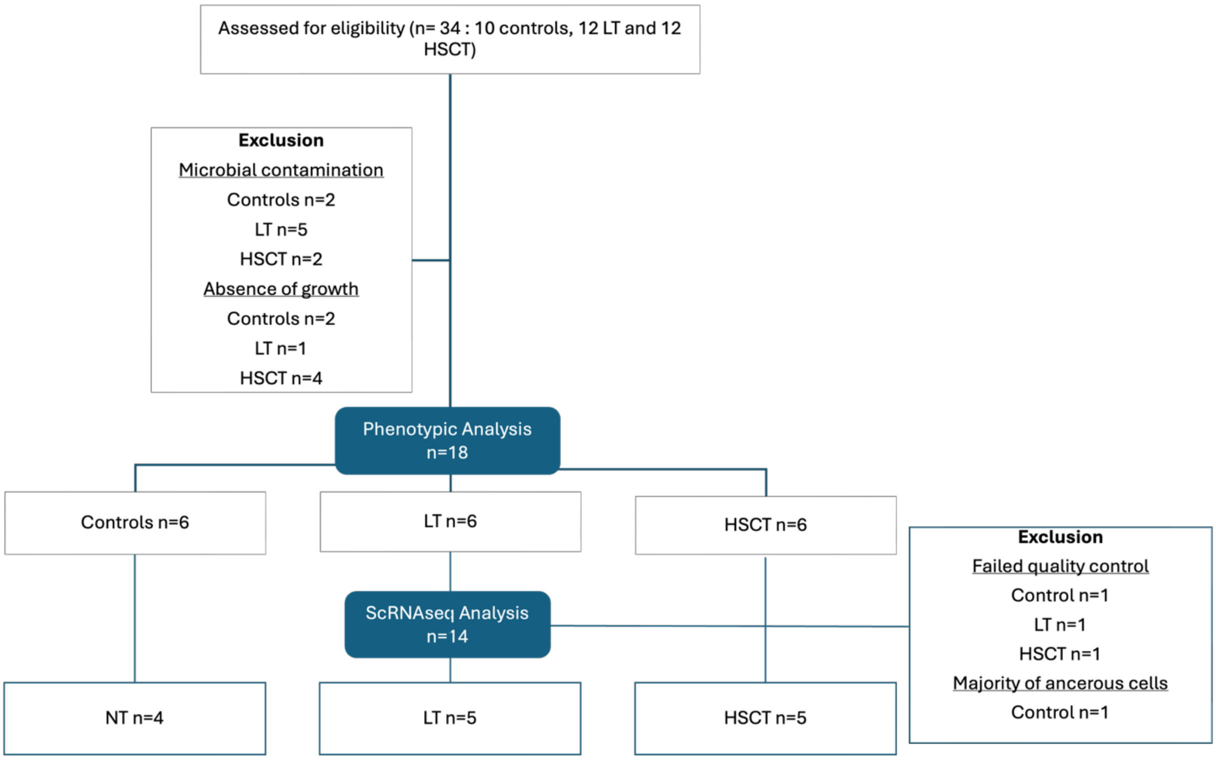


**Figure S1 – Flow chart of the study**

*Flow diagram showing patient enrollment eligibility assessment and downstream analyses. A total of 34 patients were assessed for eligibility. Nine samples were excluded du to microbial contamination and seven for failure of epithelial cultures to grow, leaving 18 patients (6 patients per group (NT, HSCT and LT)), retained for analysis. All these eighteen HAE underwent phenotypic characterization in a previous study (BioRxvi). In this study, these eighteen patient-derived HAE proceeded to scRNA-seq (4 NT, 5 HSCT and 5 LT) ; four samples were excluded du to failure to meet quality control thresholds, resulting in final dataset of fourteen samples included in the scRNA-seq analysis.*

**Table S1 – Non-transplant characteristics**

| Parameter | NT1 | NT2 | NT3 | NT4 | NT5 | NT6 |
|---|---|---|---|---|---|---|
| **Age, sex** | 67, F | 50, M | 67, M | 74, M | 62, F | 76, M |
| **History of smoking, PY** | 40 | 20 | 20 | No | No | 40 |
| **Indication for Bronchoscopy** | Tabacco nodule | COPD nodule | Tabacco nodule | Nodule adenopathy | adenopathy | Emphysema Nodule |
| **FEV1 (% predicted)** | 1610 (71) | 1610 (47) | 2100 (82) | 2330 (89) | 2510 (83) | 1560 (48) |
| **FVC (% predicted)** | 2460 (85) | 2200 (51) | 2620 (82) | 3460 (102) | 3170 (86) | 2130 (51) |
| **FEV1/FVC** | 65 | 73 | 67 | 80 | 79 | 73 |

*NT: non-transplant; PY: pack-year (number of cigarette packs per day x the number of years of smoking) FEV1: forced expiratory volume in 1 second, FVC: forced vital capacity*

**Table S2 – HSCT recipients characteristics**

| Parameters | HSCT1 | HSCT2 | HSCT3 | HSCT4 | HSCT5 | HSCT6 |
|---|---|---|---|---|---|---|
| **Age, sex** | 22, F | 51, F | 59, F | 48, M | 30, M | 55, F |
| **Donor age, sex** | 21, M | 57, F | 32, F | 23, F | 32, M | 65, F |
| **History of smoking, PY** | 0 | 0 | 26 | 20 | 0 | 0 |
| **Indication for HSCT** | AML | AML | NHL T | ALL | Sickle cell Disease | ALL |
| **Type of graft** | Non-Related donor<br>HLA 10/10<br>PSC<br>Non T dep | Non-Related donor<br>HLA 10/10<br>PSC<br>Non T dep | Haplo<br>PSC n<br>Non-T dep | Haplo<br>PSC<br>Non-T dep | Haplo<br>BM<br>Non-T dep | Non-Related donor<br>HLA 10/10<br>PSC<br>Non T dep |
| **GVHD prevention regimen** | CsA 3mg/kg/d<br>MTX 15mg/m2 | FK 0.03mg/kg/d<br>MTX 15mg/m2 | Cy 50mg/kg/d<br>FK 0.03mg/kg/d<br>MMF 15mg/kg x3/d | Cy 50mg/kg/d<br>FK 0.03mg/kg/d<br>MMF 15mg/kg x3/d | Cy 50mg/kg/d<br>FK 0.03mg/kg/d<br>MMF 15mg/kg x3/d | FK 0.03mg/kg/d<br>MTX 15mg/m2 |
| **Days post transplant to biopsy** | 210 | 843 | 343 | 1008 | 120 | 359 |
| **Serology (R/D)** | CMV -/-<br>Toxo -/-<br>EBV -/+ | CMV +/+<br>Toxo +/-<br>EBV +/+ | CMV -/-<br>Toxo +/+<br>EBV+/+ | CMV +/+<br>Toxo +/-<br>EBV+/- | CMV -/-<br>Toxo -/-<br>EBV +/+ | CMV +/+<br>Toxo +/+<br>EBV -/+ |
| **Hematopoietic chimerism at time of biopsy, % donor** | 100% | 93% | 100% | 100% | 99% | 99% |

| | | | | | | |
|---|---|---|---|---|---|---|
| **Conditioning regimen (total dose)** | MAC<br>ATG (1.5g)<br>Cy (5.6g)<br>TBI (12Gy) | MAC<br>ATG (2400mg)<br>Flu (275mg)<br>Treosulfan (77g) | RIC<br>Clofarabine (270mg)<br>Cy(2.3g)<br>Flu (270mg)<br>Melphalan (198mg) | MAC<br>ATG (700mg)<br>Etoposide (4.2g)<br>TBI (10Gy) | RIC<br>ATG (300mg)<br>Thiotepa (651mg)<br>Cy (1822mg)<br>Flu (257mg)<br>TBI (2Gy) | RIC<br>ATG (500mg)<br>Flu (275mg)<br>TBI (8Gy) |
| **GVHD** | Grade 3 skin aGVHD | Mild skin cGVHD<br>Ø at the time of biopsy | Mild skin cGVHD | Mild skin cGVHD | Severe skin cGVHD sclerodermiform | Grade 3 Digestive aGVHD |
| **History of RVI HSCT (virus, days from graft)** | HPIV3, 352, SARS-CoV-2 470 | MPV, SARS-CoV-2, RHV 441 | RHV, 151 & 263, SARS-CoV-2, 753 | HPIV3 & RHV 83 | RHV -134, HPIV3 156, & 230, RHV Cor-OC43 267, RSV 355 | None |
| **History of RBI HSCT (bacteria, days from graft)** | None | None | None | None | None | None |
| **IS drugs at biopsy** | Ruxo<br>5mgx2 | AZA | Pred 10mg<br>FK 2.5mgx2 | stopped (90 d) | Pred 5mg<br>FK 2.5mg | PONA<br>15mg/j |

*HSCT: hematopoietic stem cell transplantation; PY: pack-year (number of cigarette packs per day x the number of years of smoking); Haplo: haploidentical; T dep : T-cells depleted; PSC: peripheral stem cell; HLA: human leukocyte antigen; AML : acute myeloid leukemia; ALL: acute lymphoid leukemia; MTX: methotrexate; MMF: mycophenolate mofetil; CSA: cyclosporin; R: recipient; D: donor; CMV: cytomegalovirus; EBV: Epstein Barr virus; Toxo: toxoplasmosis; MAC: myeloablative conditioning; RIC: reduced-intensity conditioning; ATG: anti thymoglobuline, Flu: fludarabine: Cy: cyclophosphamide; TBI: total body irradiation; GVHD: graft versus host disease; RVI: respiratory viral infection; HPIV3: human parainfluenuza virus type 3; RHV:*

*rhinovirus; RSV: respiratory syncytial virus; MPV: metapneumovirus; CoroOC43: coronavirus OC43 RBI: respiratory bacterial infection; IS: immunosuppressive; FK: tacrolimus; AZA: Azacitidine; Ruxo : ruxolitinib; Pred. ; prednisone; PONA: ponatinib*

**Table S3 – Lung transplant recipients characteristics**

| Parameter | LT1 | LT2 | LT3 | LT4 | LT5 | LT6 |
|---|---|---|---|---|---|---|
| **Sex-mismtach** | No | Yes | NA | Yes | No | No |
| **History of smoking, PY of recipient** | 30 | 0 | 40 | 25 | 10 | 10 |
| **Donor age, sex** | 22 | 71 | NA | 36 | 46 | 52 |
| **History of smoking, PY of donor** | 4 | 10 | NA | 0 | 0 | 0 |
| **Indication for LT** | COPD | Emphyse ma | IPF | IPF /PAH | IPF | NSIP |
| **Type of transplantatio n** | Bi-pulm | Bi-pulm | Bi-pulm | Bi-pulm | Bi-pulm | Bi-pulm |
| **Serology (R/D)** | CMV +/-<br>EBV +/+ | CMV +/+<br>EBV +/+ | CMV -/-<br>EBV+/+ | CMV +/-<br>EBV+/- | CMV +/-<br>EBV +/+ | CMV +/+<br>EBV +/- |
| **Blood Group (R/D)** | A +/A + | A+/A+ | B+/NA | A+/A+ | O+/O+ | O+/O- |
| **Primary graft dysfunction** | No | No | No | No | No | No |
| **Days post transplant to biopsy** | 725 | 574 | 545 | 276 | 96 | 334 |
| **Ischemic duration** | 5h37 | 4h58 | NA | 10h | 4h53 | 9h45 |
| **Gastroesopha geal reflux** | Yes | No | Yes | No | No | Yes |

| | | | | | | |
|---|---|---|---|---|---|---|
| **History of RVI after Tx (virus, days from graft)** | RHV, 477 | SARS-CoV-2, 67 | SARS-CoV-2, 511 | None | None | RHV -44 ; RSV -107 |
| **History of RBI after Tx (bacteria, days from graft)** | None | None | *P. aer*, 311 | None | None | Coryne., 483 |
| **IS drug at biopsy** | FK 3.75mg x2/d<br>MMF 1g x2/d<br>Pred 5mg | FK 3mgx2/d<br>MMF 180mg x2/d<br>Pred 20mg | FK 4.5mg/d<br>MMF 360mgx2<br>Pred 7.5mg | FK 2.75mg x2/d<br>MMF 500mg x2/d<br>Pred 5mg | FK 2.5mgx2 /d<br>MMF 1g x2/d<br>Pred 22.5mg | FK 2.5mg2/ d<br>MMF 500mg x2/d<br>Pred 12.5mg |

*LT : lung transplantation; PY: pack-year (number of cigarette packs per day x the number of years of smoking); Bi-pulm: bi-pulmonary; IPF: idiopathic pulmonary fibrosis; COPD: chronic obstructive disease; PAH: pulmonary hypertension; NSIP: non-specific interstitial pneumonia; R: recipient; D: donor; CMV: cytomegalovirus; EBV: Epstein Barr virus; Toxo: toxoplasmosis; RVI: respiratory viral infection; HPIV3: human parainfluenuza virus type 3; RHV: rhinovirus; RSV: respiratory syncytial virus; RBI: respiratory bacterial infection; P. aer : pseudomonas aerugninosa ; Coryne. Corynebacterium; IS: immunosuppressive; MMF: mycophenolate mofetil; FK: tacrolimus; Pred: prednisone.*

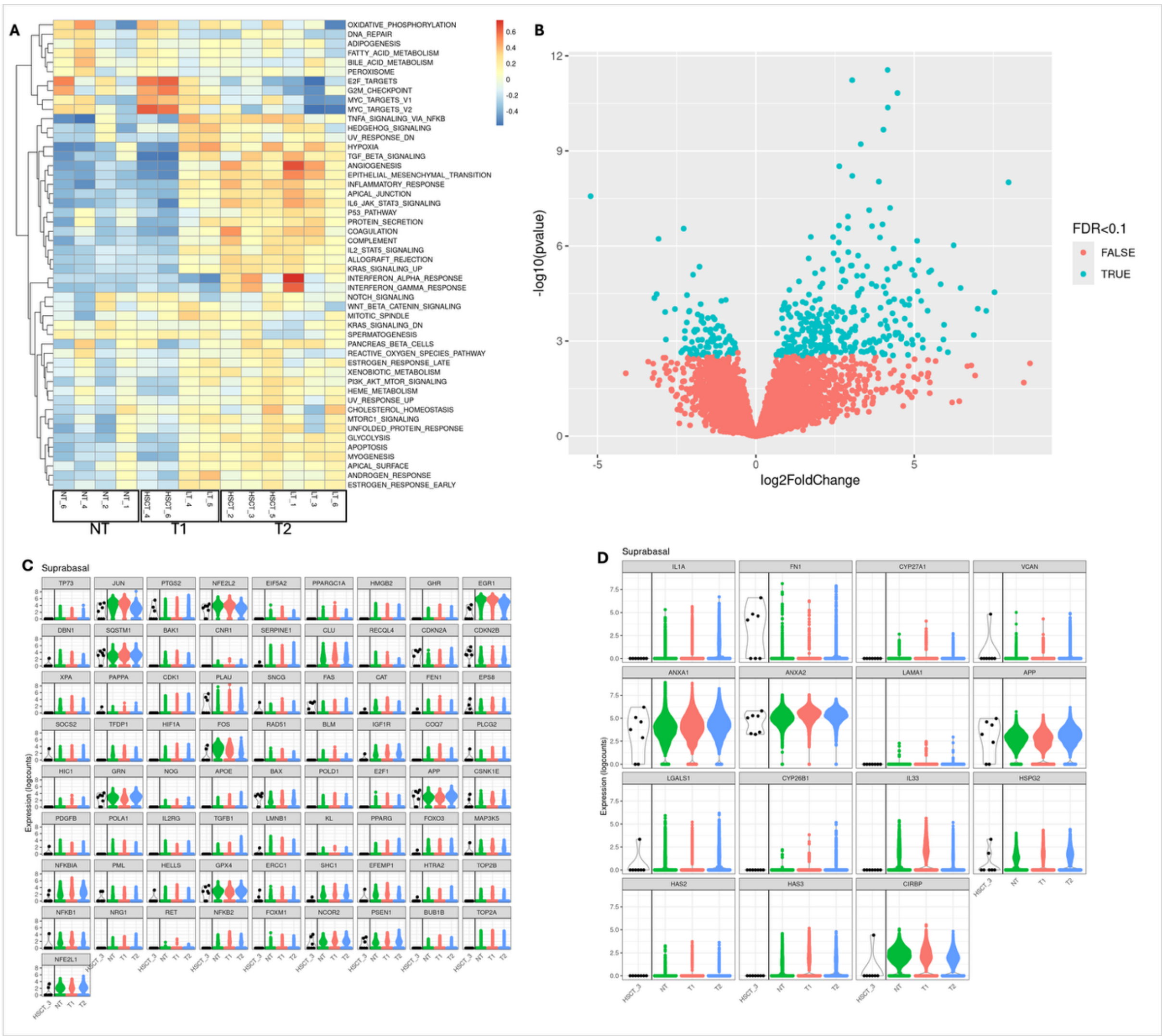


**Figure S2 - Transcriptomic profile and expression of Aging-related and DAMPs-related genes of supra basal cells**

*(A) Heatmap displaying the relative expression of 50 signaling pathway-associated gene across individual samples, with subgroups in black boxes (NT: non-transplant; T1: transplant group 1; T2: transplant group 2), within each epithelial cell population. Gene expression values are scaled by row (gene) to highlight up or down regulated patterns. Samples are arranged by hierarchical clustering to emphasize shared transcriptional signatures within suprabasal cells(A). (B) Volcano plot depicting gene expression of suprabasal cells.. Significantly differentially expressed genes are highlighted in blue (adjusted p value <0.1), with the x axis representing the log2 fold change and the y axis the -log10 (p value). (C) Violin plots display log-normalized expression levels of key aging-associated genes in suprabasal cells. Samples are grouped into three clinical categories: NT (green), T1 (orange) and T2 (blue), HSCT3, the outlier who subsequently developed BOS, is also depicted (black). Each violin represents the*

*per-cell expression distribution for the indicated epithelial subtype within each group. (B) Violin plots display log-normalized expression levels of key DAMPs-associated in suprabasal cells. Samples are grouped into three clinical categories: NT (green), T1 (orange) and T2 (blue), HSCT3, the outlier who subsequently developed BOS, is also depicted (black). Each violin represents the per-cell expression distribution for the indicated epithelial subtype within each group.*

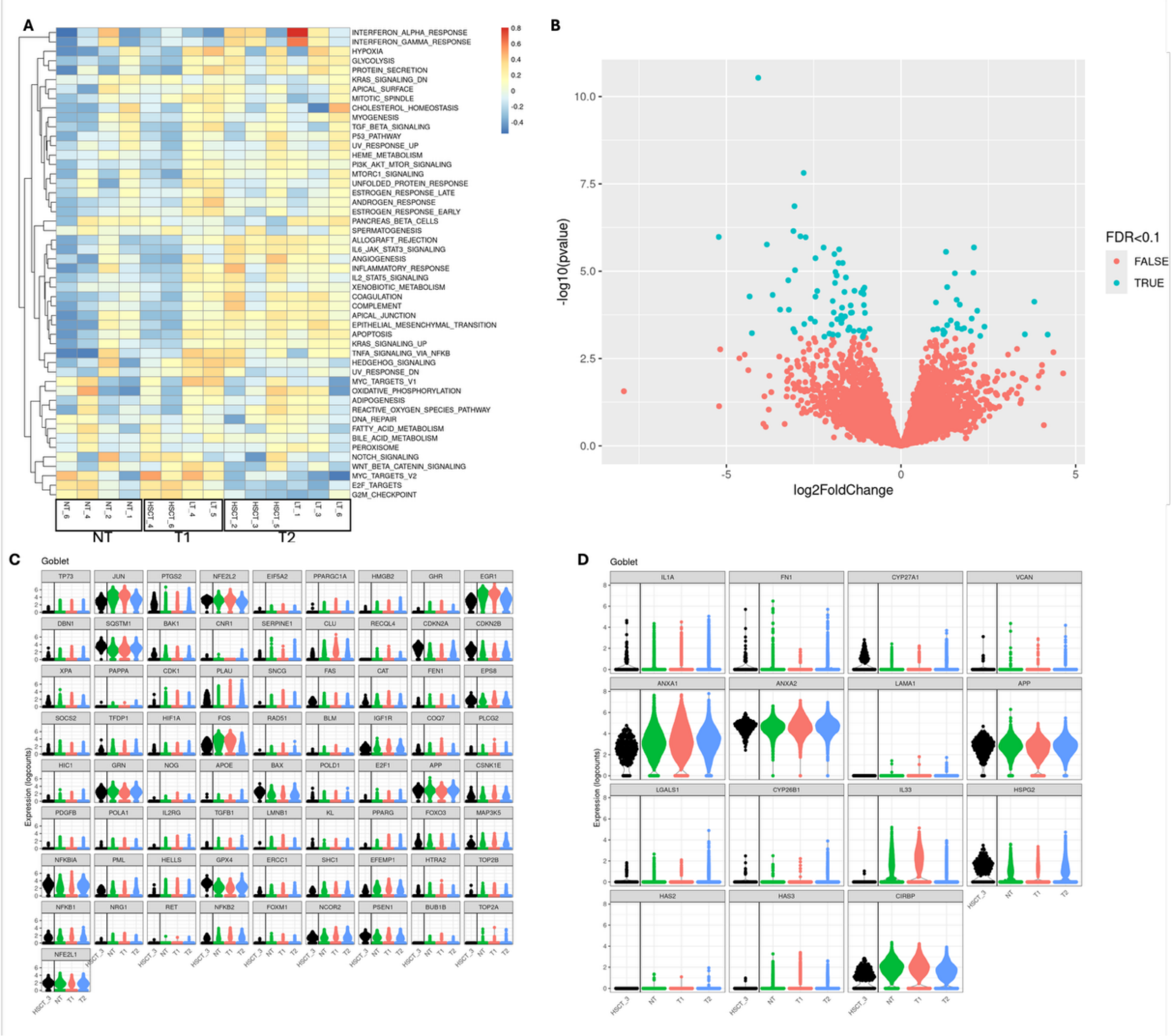


**Figure S3 - Transcriptomic profile and expression of Aging-related and DAMPs-related genes of goblet cells**

*(A) Heatmap displaying the relative expression of 50 signaling pathway-associated gene across individual samples, with subgroups in black boxes (NT: non-transplant; T1: transplant group 1; T2: transplant group 2), within each epithelial cell population. Gene expression values are scaled by row (gene) to highlight up or down regulated patterns. Samples are arranged by hierarchical clustering to emphasize shared transcriptional signatures within suprabasal*

*cells(A). (B) Volcano plot depicting gene expression of suprabasal cells. Significantly differentially expressed genes are highlighted in blue (adjusted p value <0.1), with the x axis representing the log2 fold change and the y axis the -log10 (p value). (C) Violin plots display log-normalized expression levels of key aging-associated genes in goblet cells. Samples are grouped into three clinical categories: NT (green), T1 (orange) and T2 (blue), HSCT3, the outlier who subsequently developed BOS, is also depicted (black). Each violin represents the per-cell expression distribution for the indicated epithelial subtype within each group. (B) Violin plots display log-normalized expression levels of key DAMPs-associated goblet cells. Samples are grouped into three clinical categories: NT (green), T1 (orange) and T2 (blue), HSCT3, the outlier who subsequently developed BOS, is also depicted (black). Each violin represents the per-cell expression distribution for the indicated epithelial subtype within each group.*

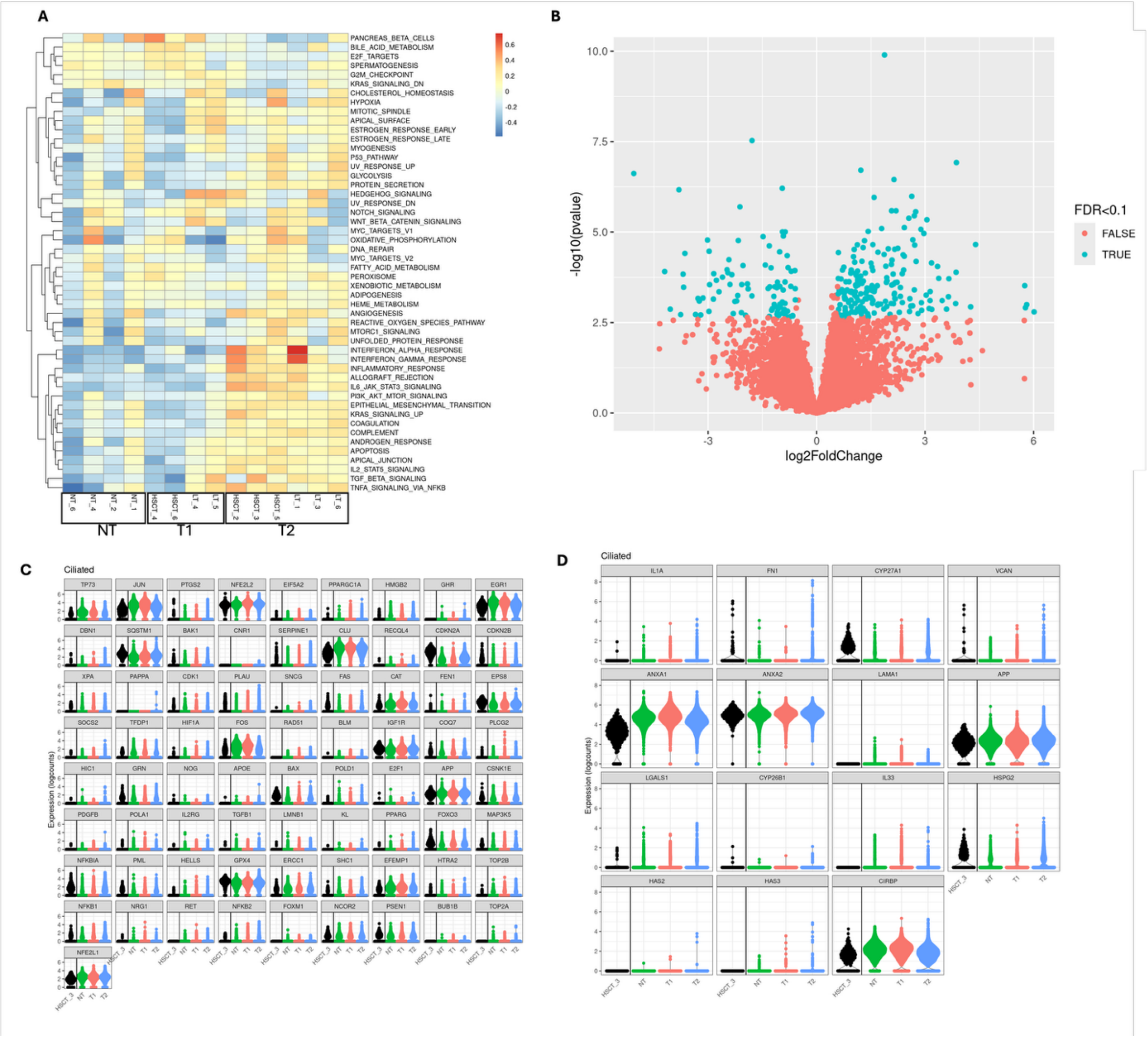

**Figure S4 - Transcriptomic profile and expression of Aging-related and DAMPs-related genes of ciliated cells**

*(A) Heatmap displaying the relative expression of 50 signaling pathway-associated gene across individual samples, with subgroups in black boxes (NT: non-transplant; T1: transplant group 1; T2: transplant group 2). Gene expression values are scaled by row (gene) to highlight up or down regulated patterns. Samples are arranged by hierarchical clustering to emphasize shared transcriptional signatures within ciliated cells(A). (B) Volcano plot depicting gene expression of ciliated cells. Significantly differentially expressed genes are highlighted in blue (adjusted p value <0.1), with the x axis representing the log2 fold change and the y axis the -log10 (p value). (C) Violin plots display log-normalized expression levels of key aging-associated genes in ciliated cells. Samples are grouped into three clinical categories: NT (green), T1 (orange) and T2 (blue), HSCT3, the outlier who subsequently developed BOS, is also depicted (black). Each violin represents the per-cell expression distribution for the indicated epithelial subtype within each group. (B) Violin plots display log-normalized expression levels of key DAMPs-associated in ciliated cells. Samples are grouped into three clinical categories: NT (green), T1 (orange) and T2 (blue), HSCT3, the outlier who subsequently developed BOS, is also depicted (black). Each violin represents the per-cell expression distribution for the indicated epithelial subtype within each group.*

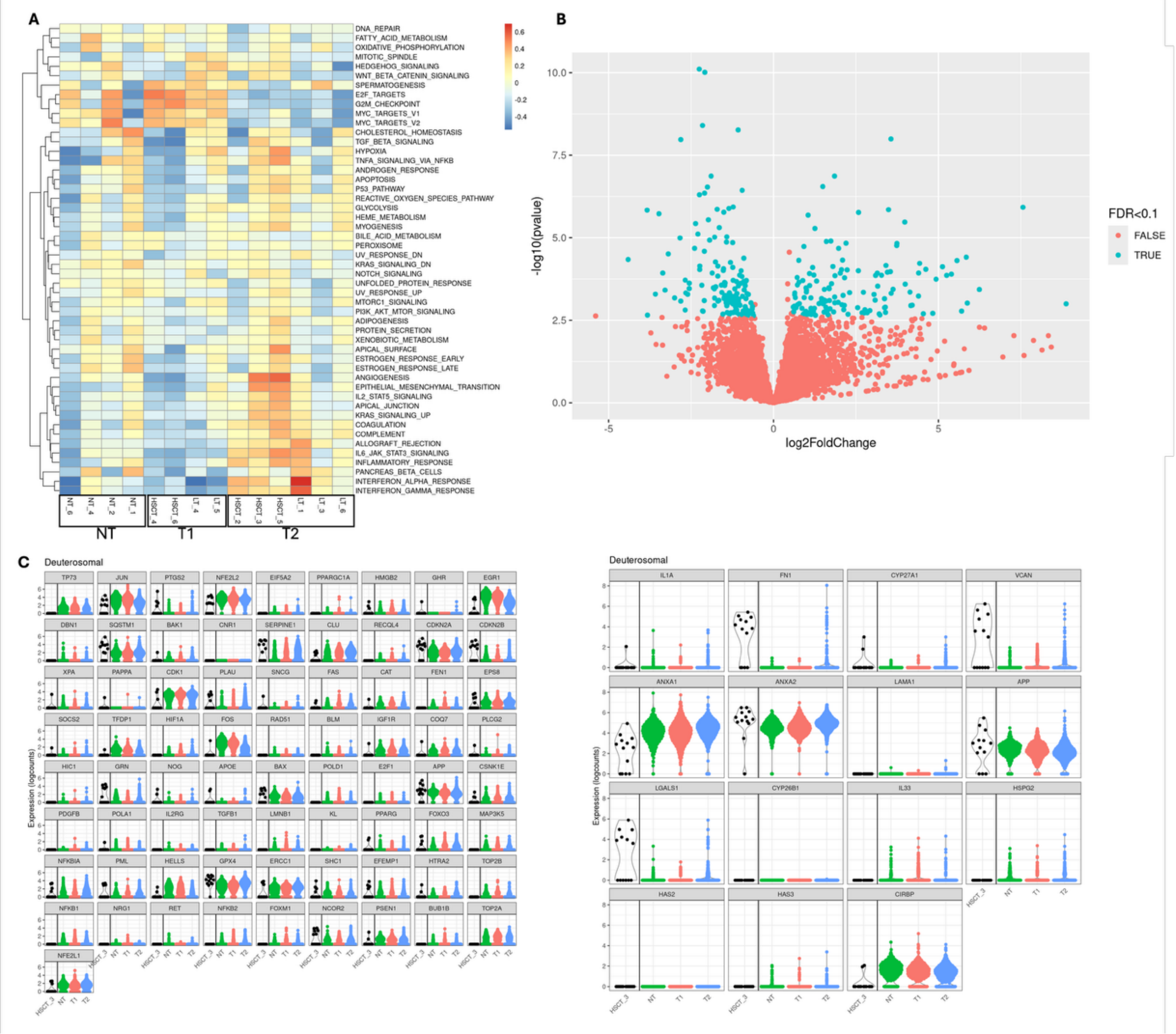

**Figure S5 - Transcriptomic profile and expression of Aging-related and DAMPs-related genes of deuterosomal cells**

*(A) Heatmap displaying the relative expression of 50 signaling pathway-associated gene across individual samples, with subgroups in black boxes (NT: non-transplant; T1: transplant group 1; T2: transplant group 2). Gene expression values are scaled by row (gene) to highlight up or down regulated patterns. Samples are arranged by hierarchical clustering to emphasize shared transcriptional signatures within deuterosomal cells(A). (B) Volcano plot depicting gene expression of deuterosomal cells. Significantly differentially expressed genes are highlighted in blue (adjusted p value <0.1), with the x axis representing the log2 fold change and the y axis the -log10 (p value). (C) Violin plots display log-normalized expression levels of key aging-associated genes in deuterosomal cells. Samples are grouped into three clinical categories: NT (green), T1 (orange) and T2 (blue), HSCT3, the outlier who subsequently developed BOS, is also depicted (black). Each violin represents the per-cell expression distribution for the indicated*

*epithelial subtype within each group. (B) Violin plots display log-normalized expression levels of key DAMPs-associated in deuterosomal cells. Samples are grouped into three clinical categories: NT (green), T1 (orange) and T2 (blue), HSCT3, the outlier who subsequently developed BOS, is also depicted (black). Each violin represents the per-cell expression distribution for the indicated epithelial subtype within each group.*

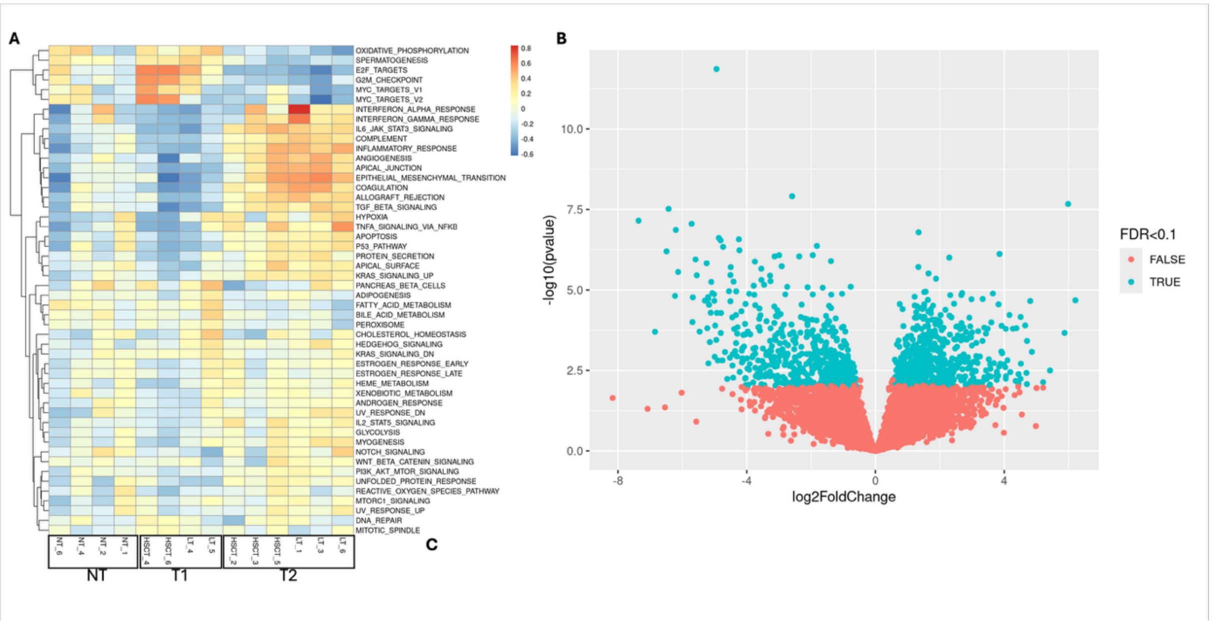


**Figure S6 - Transcriptomic profile of rare cells**

*(A) Heatmap displaying the relative expression of 50 signaling pathway-associated gene across individual samples, with subgroups in black boxes (NT: non-transplant; T1: transplant group 1; T2: transplant group 2). Gene expression values are scaled by row (gene) to highlight up or down regulated patterns. Samples are arranged by hierarchical clustering to emphasize shared transcriptional signatures within rare cells(A). (B) Volcano plot depicting gene expression of rare cells. Significantly differentially expressed genes are highlighted in blue (adjusted p value <0.1), with the x axis representing the log2 fold change and the y axis the -log10 (p value).*

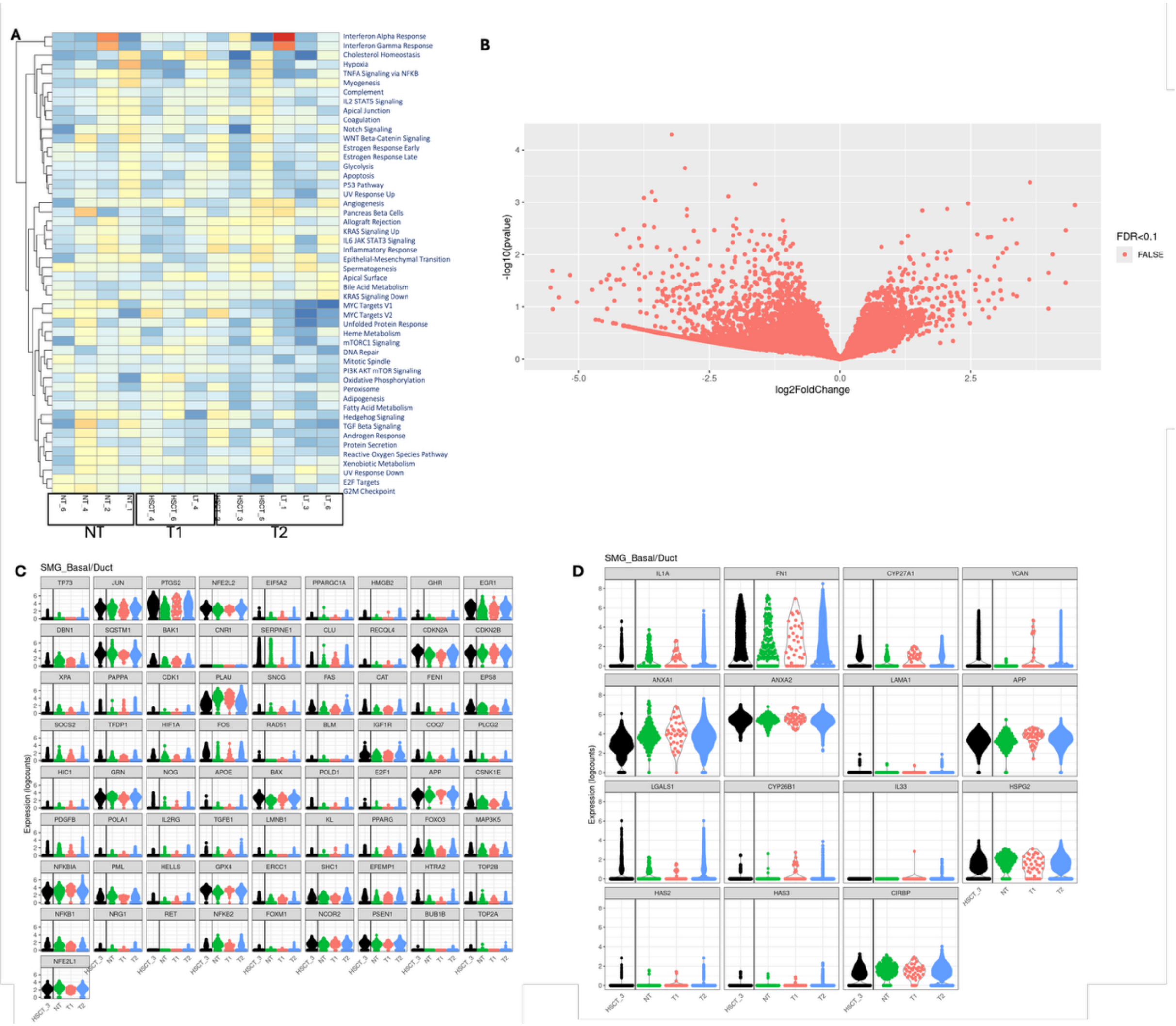


**Figure S7 - Transcriptomic profile and expression of Aging-related and DAMPs-related genes of SMG basal duct cells**

*(A) Heatmap displaying the relative expression of 50 signaling pathway-associated gene across individual samples, with subgroups in black boxes (NT: non-transplant; T1: transplant group 1; T2: transplant group 2). Gene expression values are scaled by row (gene) to highlight up or down regulated patterns. Samples are arranged by hierarchical clustering to emphasize shared transcriptional signatures within SMG basal duct cells(A). (B) Volcano plot depicting gene expression of SMG basal duct cells. Significantly differentially expressed genes are highlighted in blue (adjusted p value <0.1), with the x axis representing the log2 fold change and the y axis the -log10 (p value). (C) Violin plots display log-normalized expression levels of key aging-associated genes in SMG basal duct cells. Samples are grouped into three clinical categories: NT (green), T1 (orange) and T2 (blue), HSCT3, the outlier who subsequently developed BOS, is*

*also depicted (black). Each violin represents the per-cell expression distribution for the indicated epithelial subtype within each group. (B) Violin plots display log-normalized expression levels of key DAMPs-associated in SMG basal duct cells. Samples are grouped into three clinical categories: NT (green), T1 (orange) and T2 (blue), HSCT3, the outlier who subsequently developed BOS, is also depicted (black). Each violin represents the per-cell expression distribution for the indicated epithelial subtype within each group.*

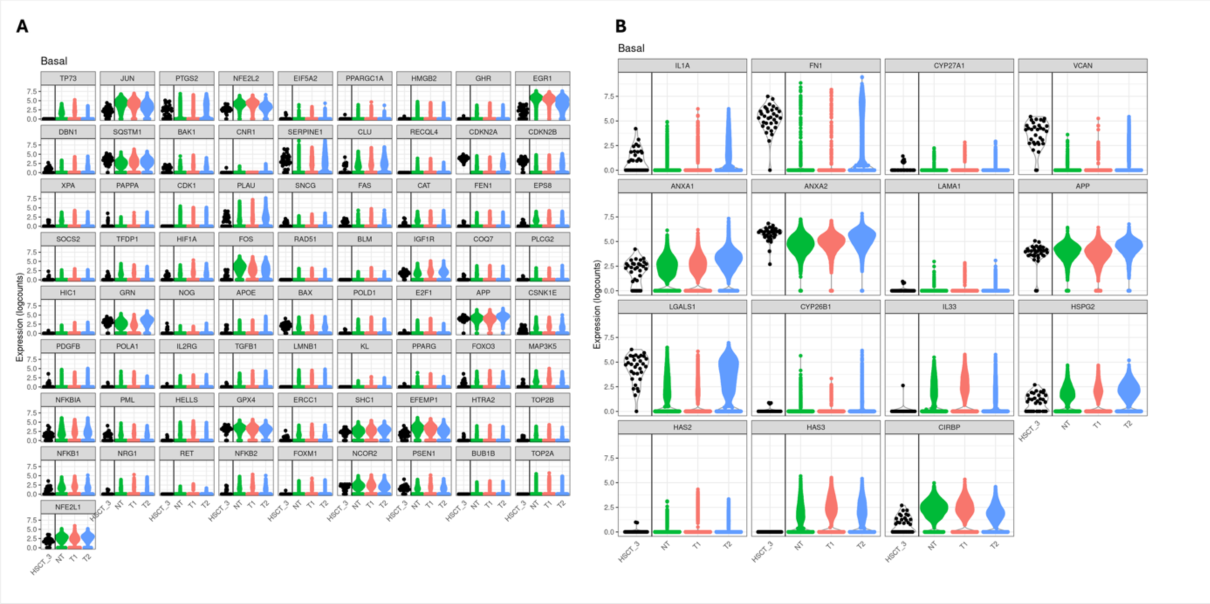


**Figure S8 – Expression of Aging-related and DAMPs-related genes in basal cells**

*(A) Violin plots display log-normalized expression levels of key aging-associated genes in basal cells. Samples are grouped into three clinical categories: NT (green), T1 (orange) and T2 (blue), HSCT3, the outlier who subsequently developed BOS, is also depicted (black). Each violin represents the per-cell expression distribution for the indicated epithelial subtype within each group. (B) Violin plots display log-normalized expression levels of key DAMPs-associated in basal cells. Samples are grouped into three clinical categories: NT (green), T1 (orange) and T2 (blue), HSCT3, the outlier who subsequently developed BOS, is also depicted (black). Each violin represents the per-cell expression distribution for the indicated epithelial subtype within each group.*

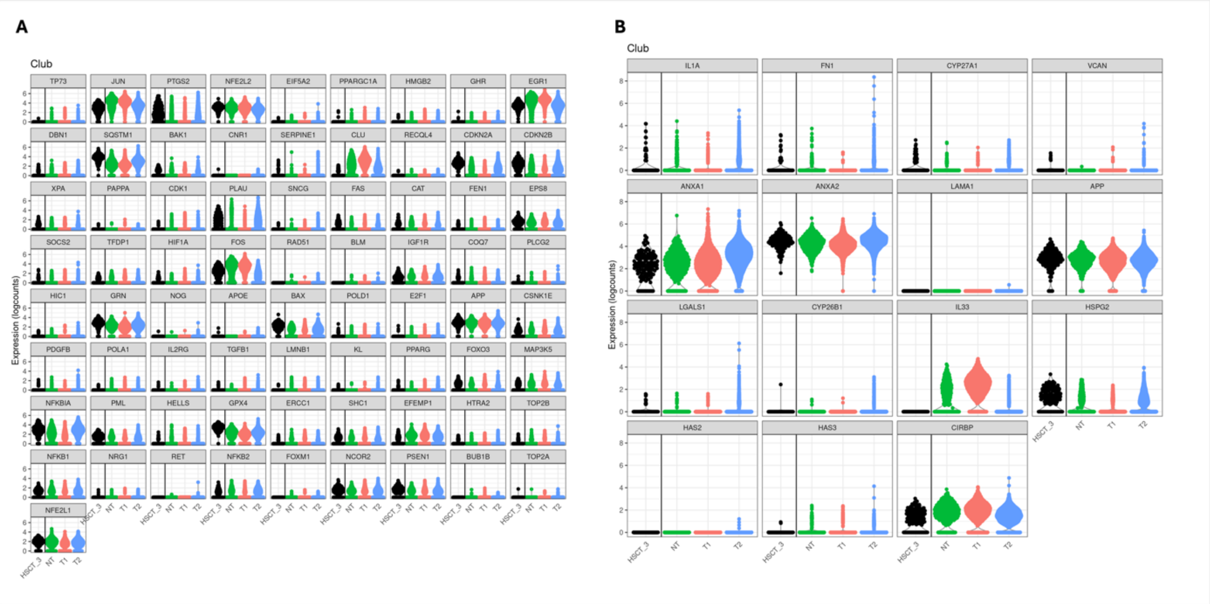


**Figure S9 – Expression of Aging-related and DAMPs-related genes in club cells**

*(A) Violin plots display log-normalized expression levels of key aging-associated genes in club cells. Samples are grouped into three clinical categories: NT (green), T1 (orange) and T2 (blue), HSCT3, the outlier who subsequently developed BOS, is also depicted (black). Each violin represents the per-cell expression distribution for the indicated epithelial subtype within each group. (B) Violin plots display log-normalized expression levels of key DAMPs-associated in club cells. Samples are grouped into three clinical categories: NT (green), T1 (orange) and T2 (blue), HSCT3, the outlier who subsequently developed BOS, is also depicted (black). Each violin represents the per-cell expression distribution for the indicated epithelial subtype within each group.*
